# Phosphorylation-dependent tuning of mRNA deadenylation rates

**DOI:** 10.1101/2024.10.18.618793

**Authors:** James A.W. Stowell, Conny W.H. Yu, Zhuo A. Chen, Giselle Lee, Tomos Morgan, Ludwig Sinn, Sylvie Agnello, Francis J. O’Reilly, Juri Rappsilber, Stefan M.V. Freund, Lori A. Passmore

**Affiliations:** MRC Laboratory of Molecular Biology, Cambridge CB2 0QH UK; Technische Universität Berlin, Chair of Bioanalytics, 10623 Berlin, Germany; Wellcome Centre for Cell Biology, University of Edinburgh, EH9 3BF, UK

**Keywords:** exonuclease, gene expression, intrinsically disordered proteins, phosphorylation, mRNA decay

## Abstract

mRNA decay is a major determinant of gene regulation that is controlled through shortening of mRNA poly(A) tails by the Ccr4-Not complex. The specificity of deadenylation can be mediated through RNA adaptors – RNA-binding proteins that tether substrate mRNAs to Ccr4-Not in a regulated and context-specific manner. Interaction with Ccr4-Not is mediated by intrinsically disordered regions (IDRs) within the RNA adaptors. Due to the difficulty in studying large IDR-containing complexes, the determinants of specificity and their regulation remain unclear. Here we use structural biology and biochemical reconstitution to show that dispersed segments within IDRs of RNA adaptors bind to several distinct binding sites on Ccr4-Not through multivalent interactions. We further demonstrate that binding can be modulated by phosphorylation, altering the consequent deadenylation rate in a continuously tunable manner. This mechanism is broadly applicable in evolutionarily divergent IDRs from multiple RNA adaptors including fission yeast Puf3, and human Pumilio/PUM1 and Tristetraprolin/TTP. Together, our work suggests that multivalent interactions and phosphorylation represent conserved strategies for regulating gene expression. Thus, in response to cellular cues, mRNA decay can be regulated by a graded mechanism, rather than a bistable on/off switch, rationalizing how post-transcriptional gene expression is fine-tuned.

---

Regulation of mRNA 3’-poly(A) tail length is critical for the control of mRNA half-life and translational efficiency ^1^. Shortening of poly(A) tails is mediated by Ccr4-Not and Pan2-Pan3, highly conserved multi-protein complexes that contain exonucleases termed deadenylases ^2^. Ccr4-Not contains seven core subunits including the deadenylase enzymes Ccr4/CNOT6/ CNOT6L and Caf1/CNOT7/CNOT8, which assemble around the ∼200 kDa Not1/CNOT1 scaffold (Extended Data Fig. 1A) ^3^.

Ccr4-Not is targeted to specific mRNAs through a diverse range of RNA adaptor proteins that respond to cellular signals, developmental cues and the translational state of the ribosome^1^. RNA adaptor proteins commonly have a modular architecture, with a structurally conserved RNA-binding domain that interacts with specific cis-acting RNA sequence elements, as well as intrinsically disordered regions (IDRs) that recruit Ccr4-Not. RNA adaptors therefore tether specific transcripts to the deadenylation machinery to accelerate poly(A) tail shortening ^4-8^. In this ‘tethering model’, short linear motifs (SLiMs) bind to one of several hydrophobic pockets located on the Not1, Not3/CNOT3 and Not9/Caf40/Rcd1/CNOT9 subunits (Extended Data Fig. 1A) ^9^. For example, distinct SLiMs in the RNA-binding proteins UnKempt and Roquin interact with both CNOT9 and the C-terminal NOT module (a structured region of Not1, Not2 and Not3) ^10,11^.

Disruption of SLiMs in RNA adaptors leads to a reduced ability to bind Ccr4-Not, thereby decreasing RNA turnover. However, a single SLiM is often insufficient to account for the full repressive activity of the IDR, both *in vitro* and *in vivo*. For example, the AU-rich element (ARE) binding protein Tristetraprolin (TTP or ZFP36) interacts with an N-terminal HEAT repeat in CNOT1 through a highly conserved RLP(ϕ)F SLiM (where ϕ is a hydrophobic residue) ^12^.

Deletion of this motif only partially stabilizes the TTP target pro-inflammatory mRNAs *in vivo*, and does not have the severe autoimmune phenotype of knockout mice ^13^. Thus, it is likely that additional elements within TTP contribute to its binding to Ccr4-Not and stimulation of deadenylation.

Discrete SLiMs have been difficult to identify in some RNA adaptors. For example, Drosophila Pumilio and human PUM1 contain long IDRs with multiple regions that can repress reporter mRNAs via interaction with Ccr4-Not ^14-17^. The large size, low sequence conservation and lack of structure in these IDRs generate challenges in studying their binding mechanisms. Thus, despite multiple lines of evidence supporting the essentiality of IDRs and SLiMs in RNA turnover, mechanistic details are lacking.

Here, we use *in vitro* reconstitution and structural biology to address how RNA adaptors bind and stimulate Ccr4-Not. We find that they act via a multipartite IDR binding mechanism that is tuned by phosphorylation. This is reminiscent of a mechanism used by transcription factors to regulate mRNA synthesis ^18,19^, suggesting commonalities across the regulation of gene expression.

## Multiple regions in the Puf3 IDR are required for stimulation of Ccr4-Not

To understand how the specificity of mRNA deadenylation is mediated, we first investigated how the *S. pombe* Pumilio RNA adaptor Puf3 binds Ccr4-Not. Puf3 promotes targeted deadenylation in a reconstituted *in vitro* assay with Ccr4-Not ^20^. The first 258 residues of the N-terminal IDR in Puf3 are necessary for efficient stimulation of deadenylation (Fig. 1A-B). N and C-terminal truncations of the IDR lead to progressive decreases in activity (compare boxed regions on Fig. 1B).

**Fig. 1.**
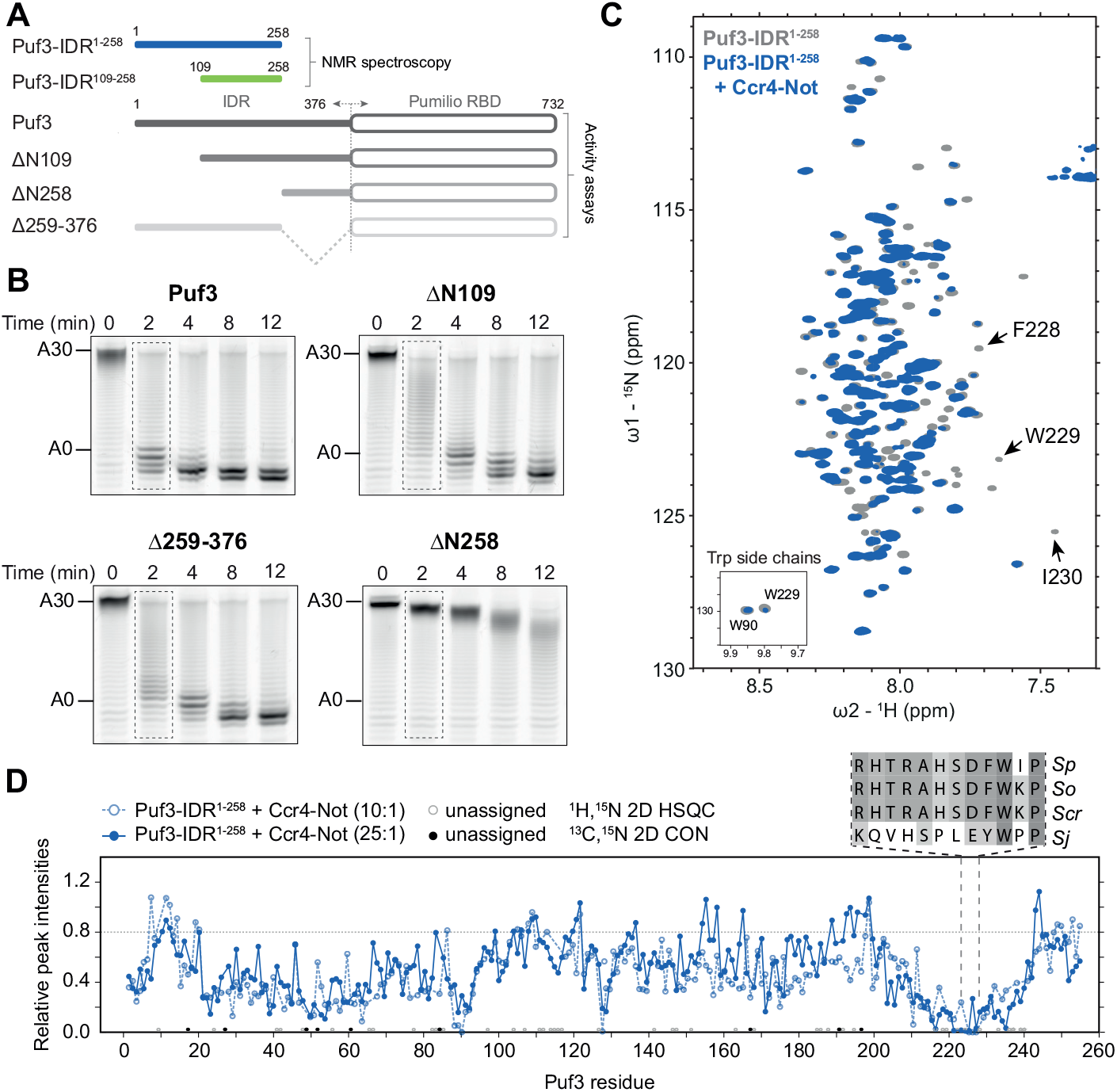
An N-terminal IDR of Puf3 is required for stimulation of deadenylation by Ccr4-Not. (**A**) Domain diagrams of *S. pombe* Puf3 constructs used in this study. Puf3 contains a 376 amino acid N-terminal IDR and a C-terminal Pumilio RNA-binding domain (RBD). Puf3-IDR^1-258^ (blue) and Puf3-IDR^109-258^ (green) NMR constructs are indicated above the full-length protein. Truncation constructs used for deadenylation assays are indicated below. (**B**) Deadenylation assays using recombinant *S. pombe* Ccr4-Not complex, MBP-tagged full-length or truncated Puf3 constructs, and 5’ fluorescently-labelled substrate RNA containing a PUF-recognition element (PRE) upstream of a 30-nt poly(A) tail. The reaction mixture was resolved using denaturing PAGE at the indicated time points. The sizes of substrate RNA with a 30 nt poly(A) tail (A30) or no poly(A) tail (A0) are indicated. (**C**) ^1^H,^15^N 2D-HSQC of 75 μM Puf3-IDR^1-258^ with (blue) or without (grey) 7.5 μM Ccr4-Not. The addition of Ccr4-Not resulted in differential line broadening across the sequence, particularly pronounced within a region spanning residues 227-231. Hydrophobic residues in this motif are highlighted in the spectra. Inset shows the side-chain resonances of tryptophans. (**D**) Relative peak intensities of Puf3-IDR^1-258^ from ^1^H,^15^N 2D-HSQC experiments in the presence of Ccr4-Not at a 10:1 ratio (light blue dotted line; from panel C) and from ^13^C-detected 2D CON experiments with Ccr4-Not at 25:1 (blue solid line; from Extended Data Fig. 2A), each compared to Puf3-IDR^1-258^ alone. Conservation of the DFW motif in *Schizosaccharomycetes* is shown in the sequence alignment. *Sp, Schizosaccharomyces pombe; So, S. octosporus*; *Scr, S. cryophilus; Sj, S. japonicus*.

Next, we used solution NMR spectroscopy to gain structural insights into the Puf3 IDR. We assigned the backbone resonances of the 258-residue Puf3 IDR, hereafter referred to as Puf3-IDR^1-258^, using a combination of ^1^H- and ^13^C-detected experiments (Fig. 1C and Extended Data Fig. 2A). A shorter construct spanning residues 109-258 (Puf3-IDR^109-258^) was used to resolve the more crowded regions of the spectra (Extended Data Fig. 2B-C), allowing us to obtain a near-complete assignment (249 out of 258 backbone resonances). Overall, the narrow dispersion of ^1^H chemical shifts in the ^1^H,^15^N 2D HSQC spectrum shows that Puf3-IDR^1-258^ is largely disordered in solution and lacks any substantial structured domains (Fig. 1C and Extended Data Fig. 2D).

To characterize the interaction between Ccr4-Not and the Puf3 IDR, we titrated the full-length 0.5-MDa unlabeled Ccr4-Not complex into isotopically labelled Puf3-IDR^1-258^. Differential line broadening (Fig. 1C, Extended Data Fig. 2A-B and 3A) was observed in the spectra, indicating that the Puf3-IDR^1-258^ intearcts with Ccr4-Not. We mapped relative peak intensity changes in spectra onto the sequence of Puf3 (Fig. 1D and Extended Data Fig. 3B-E). This revealed differential line broadening across the IDR, suggesting extensive regions of the Puf3 IDR are involved in the interaction with Ccr4-Not.

Notably, a conserved tryptophan-containing motif (residues 227-231, DFWIP in *S. pombe*, henceforth referred to as the DFW motif; Extended Data Fig. 4) shows substantial signal attenuation (Fig. 1C-D and Extended Data Fig. 3) and may represent a Ccr4-Not binding motif. Previously characterized SLiMs that bind Ccr4-Not also contain aromatic and hydrophobic residues. For example, tryptophan-containing motifs in GW182 proteins are important for interaction with CNOT1 ^21,22^.

To test the functional role of the Puf3 DFW motif, we used targeted deadenylation assays with Ccr4-Not. A purified Puf3 variant in which residues 217-232 were replaced with a Gly-Ser-Asn linker had an ∼2-fold reduced ability to stimulate Ccr4-Not mediated deadenylation compared to wild-type Puf3 (Extended Data Fig. 5A-C.) This is consistent with a role for the DFW motif in the interaction with Ccr4-Not. Still, deletion of the entire N-terminal IDR (Puf3^ΔN258^) reduces deadenylation to a much greater extent (compare to Fig. 1B) and suggests additional regions within the IDR contribute to the interaction with Ccr4-Not.

Next, we identified additional Puf3 residues that are line broadened in our NMR titration experiments and are also conserved within *Schizosaccharomyces*: L43, F65, L130 and Y176. We mutated each of these residues, along with W229, to asparagine and assessed their relative contributions to the stimulation of Ccr4-Not activity. Each individual mutation resulted in modest decreases in targeted deadenylation by Ccr4-Not (Extended Data Fig. 5D). However, combinations of these IDR point mutations have an additive effect and a protein containing all five mutations (L43N, F65N, L130N, Y176N and W229N) has a ∼3-fold decrease in deadenylation activity (Fig. 2A and Extended Data Fig. 5C). These data further support a model where extensive regions of the Puf3 IDR, including the DFW motif, are required for interaction with Ccr4-Not.

**Fig. 2.**
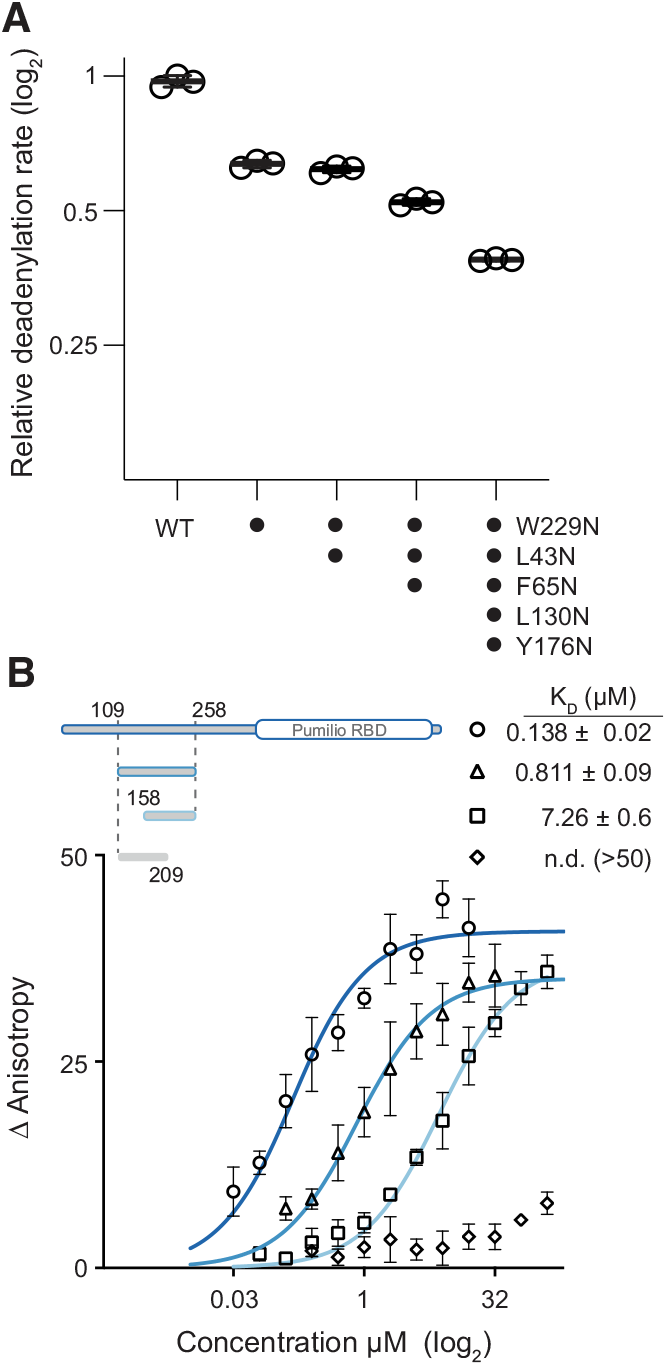
Multiple hydrophobic regions of the Puf3 IDR interact with Ccr4-Not. (**A**) Relative deadenylation rates of Ccr4-Not with mutant Puf3 proteins, normalized to Ccr4-Not activity with wild-type (WT) Puf3 (Extended Data Fig. 5A). Experiments were performed in triplicate (open circles) and averaged (line), with error bars showing the standard deviation. (**B**) Fluorescence anisotropy equilibrium analysis of Puf3 binding to labelled Ccr4-Not. Increasing concentrations of the indicated Puf3 constructs were titrated into 25 nM *S. pombe* Ccr4-Not containing a FAM-label on the Not9 subunit. Binding isotherms were fitted with a quadratic model taking account of bound ligand. Symbols represent the mean of 3 replicates with error bars showing standard deviation. Calculated Kd values (± standard error) are shown above.

To quantitate the roles of different regions of Puf3 on binding to Ccr4-Not, we prepared a fluorescently labelled Ccr4-Not complex and measured the equilibrium binding constants for Puf3 IDR truncation constructs using fluorescence anisotropy. A construct spanning residues 158-258 including the DFW motif binds to Ccr4-Not with a Kd of ∼7 μM whereas a construct spanning residues 109-202, excluding this motif, binds with substantially lower affinity (Kd > 50 μM) (Fig. 2B). A construct including both regions (residues 109-258) has a Kd of 0.8 μM, closer to the Kd of the full-length protein (0.1 μM; Fig. 2B). Together, these binding data along with NMR and mutational analyses are consistent with multiple hydrophobic motifs across the Puf3 IDR contributing to the interaction with Ccr4-Not to stimulate deadenylation activity.

## The Puf3 IDR binds multiple binding sites on Ccr4-Not

To gain insight into which surfaces of Ccr4-Not interact with Puf3, we performed crosslinking mass spectrometry (CLMS) analysis of Ccr4-Not alone and bound to Puf3 (Fig. 3A and Extended Data Fig. 6A-B). Many of the crosslinks between Puf3 and Ccr4-Not map to the Not9 subunit, but there are also crosslinks to other regions including the NOT module. Not9 is known to interact with several other RNA adaptor protein IDRs via two general mechanisms: a peptide-binding groove and two tryptophan-binding pockets ^10,21,22^. Most of the crosslinks between Puf3 and Not9 in our CLMS data locate in two clusters near the Not9 peptide-binding groove (Extended Data Fig. 6C-D). Thus, the central part of the Puf3 IDR is near Not9 and may bind directly to the peptide-binding groove.

**Fig. 3.**
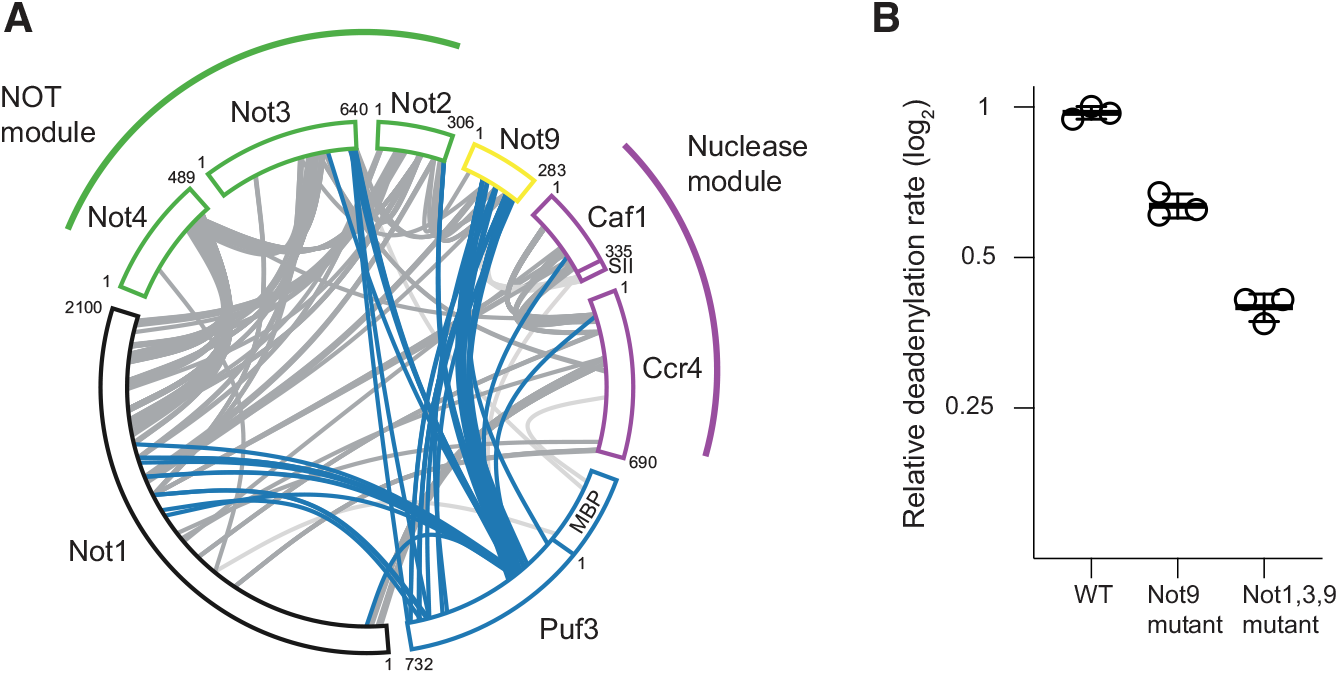
Puf3 interacts with multiple binding sites on Ccr4-Not. **(A)** Sulfo-NHS-diazirine based crosslinking mass spectrometry analysis of Ccr4-Not bound to Puf3. Blue highlighted crosslinks show interprotein crosslinks between Puf3 and Ccr4-Not subunits. Other intersubunit crosslinks are shown in grey. Subunits are colored and arranged according to the subcomplex architecture of Ccr4-Not with N- and C-terminal residue numbers for each subunit labelled. MBP, maltose-binding protein; SII, twin strep tag. **(B)** Relative deadenylation activities of wild-type (WT) Ccr4-Not and complexes containing mutations in IDR-binding pockets with WT Puf3. The mutations do not substantially affect the activity of Ccr4-Not without Puf3 (Extended Data Fig. 8). Assays were quantified as in Fig. 2A.

We used AlphaFold2 to predict the structure of a complex between Not9 and Puf3. This reproducibly places the Puf3 IDR into the peptide binding groove and the Trp-binding pockets of Not9 (Extended Data Fig. 7A-C). We examined the amino acid characteristics and evolutionary conservation of these binding sites and found that many interacting residues in Not9 are charged or hydrophobic and are invariant throughout eukaryotes (Extended Data Fig. 7D-I). We therefore hypothesized that Puf3 binds both the peptide-binding groove and the Trp-binding pockets.

To test this prediction we designed point mutations in surface residues that would disrupt the conserved interaction sites on Not9 (Extended Data Fig. 7F,I), incorporated them into Ccr4-Not and assayed the ability of Puf3 to stimulate deadenylation activity. Mutation of the Not9 binding sites reduces the ability of Ccr4-Not to accelerate deadenylation (∼1.5-2 fold reduction) (Fig. 3B and Extended Data Fig. 8). Nevertheless, this does not account for the full repressive activity of the N-terminal IDR of Puf3 (compare to Fig. 1B) suggesting that other binding sites on Ccr4-Not are also important for maximal acceleration of deadenylation by Puf3.

We next examined the evolutionary conservation of other known IDR binding sites from crystal structures of Ccr4-Not subunits in complex with SLiMs. Binding sites within Not1 and Not3 showed high conservation throughout eukaryotes, specifically in solvent exposed hydrophobic and aromatic residues (Extended Data Fig. 9). We hypothesized that these conserved binding sites could also contribute to the interaction between Puf3 and Ccr4-Not, consistent with observed crosslinks between Not3 and Puf3. Therefore, we introduced point mutations to disrupt these residues and combined these with our Not9 mutants in a fully assembled recombinant mutant Ccr4-Not complex. The Not1 and Not3 mutations further reduce the ability of Puf3 to stimulate deadenylation (Fig. 3B and Extended Data Fig. 8), consistent with a contribution of these residues towards interaction with Puf3.

Together, these data support a mode of interaction where multiple SLiMs across the Puf3 IDR bind to multiple sites on the Ccr4-Not complex, and these additive interactions are required for the full stimulatory activity of Puf3.

## Sequential phosphorylation of Puf3 tunes its ability to stimulate Ccr4-Not

IDRs are often substrates for post-translational modifications (PTMs), and these can be critical for their cellular function ^23^. The Puf3 IDR is highly enriched in serine and threonine residues, which comprise ∼30% of all residues, and some of these are phosphorylated *in vivo* (Extended Data Fig. 10A) ^24^. In *Saccharomyces cerevisiae*, Puf3p is phosphorylated in a glucose responsive manner by multiple kinases including the AGC-family kinase PKA and Sch9p ^25^. This stabilizes Puf3p-bound transcripts but does not affect RNA binding.

To test whether the *S. pombe* Puf3 IDR is similarly phosphorylated, we incubated Puf3-IDR^1-258^ with the purified AGC-family kinase Sck1 and the Sck1-activating kinase Pdk1, and analyzed the resultant protein by NMR. The backbone ^1^H resonance peaks of selected serine and threonine residues of Puf3 were shifted downfield, characteristic of phosphorylation^26^ (Fig. 4A). In total, we observed nine new resonances between 8.6-8.8 ppm and we assigned five of these to phospho-serine (pS42, pS156, pS207, pS226 and pS241) (Extended Data Fig. 10B).

**Fig. 4.**
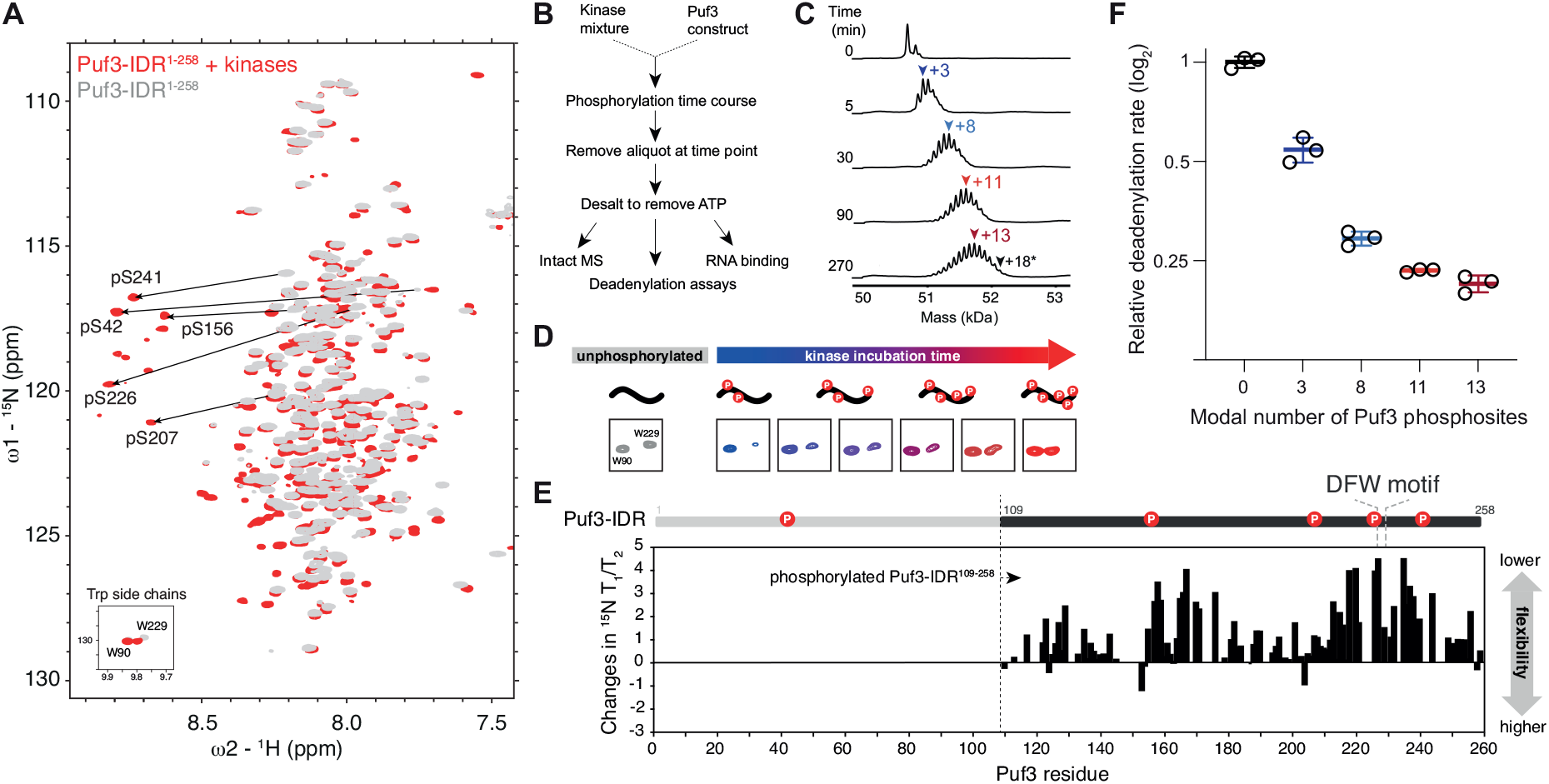
Phosphorylation of the Puf3 IDR tunes Ccr4-Not mediated deadenylation. (**A**)^1^H,^15^N 2D-HSQC spectral overlay of 75 μM unphosphorylated (grey) and phosphorylated (red) Puf3-IDR^1-258^. Puf3-IDR^1-258^ was phosphorylated using a mixture of recombinant Sck1 and Pdk1 at a 1:5 molar ratio. Substantial chemical shift perturbations are observed upon phosphorylation, particularly backbone resonances of phosphorylated serines (black arrows). Inset shows the side-chain resonances of tryptophans. (**B**) Scheme for producing differentially phosphorylated Puf3 for use in downstream experiments. (**C**) Intact mass spectrometry analysis of samples taken during the Puf3-IDR^1-376^ phosphorylation time course. Deconvoluted spectra are shown with masses consistent with modal phosphorylation states shown above each time point (colored arrows). The black arrow (+18*) at 270 min shows maximum incorporation of 18 phosphates under these conditions, with a modal number of 13 phosphorylation sites (dark red arrow). (**D**) Trp indole peaks from ^1^H,^15^N 2D-HSQC showing increasing chemical shift perturbation (CSP) for the Trp229 side-chain resonance throughout the phosphorylation time course. These data suggest that structural changes occur in the DFW motif upon phosphorylation. (**E**) Changes in ^15^N T1/T2 backbone dynamics of Puf3-IDR^109-258^ upon phosphorylation. This construct contains four out of five phosphorylation sites identified by NMR (schematic above plot). The increase in ^15^N T1/T2 indicates a decrease in backbone dynamics upon phosphorylation. Notably, the DFW motif important for Ccr4-Not binding is within a region of increased rigidity. (**F**) Relative deadenylation rates of Ccr4-Not with differentially phosphorylated full-length Puf3 proteins prepared according to panel B. IDR phosphorylation states were determined using intact-MS, indicated as modal states (see panel C). Experiments were performed in triplicate (open circles) and averaged (line), with error bars showing the standard deviation.

To gain more insight into the phosphorylation kinetics of Puf3, we performed time-resolved NMR experiments with Puf3-IDR^1-258^ and limiting concentrations of Sck1 and Pdk1 kinases. This showed sequential phosphorylation, with S42 and S156 being phosphorylated before S207, S226 and S241 (Extended Data Fig. 10C). We also analyzed phosphorylation of full-length Puf3-IDR^1-376^ over time using intact mass spectroscopy (Fig. 4B-C). This also demonstrates that Puf3-IDR^1-376^ is phosphorylated progressively: 2-4 phosphates are added in the first 5 min, and up to 18 phosphorylation sites appear over the next 4.5 h (Fig. 4C). Thus Puf3 can be differentially phosphorylated under these conditions with an increasing number of sites incorporated over time.

The resonance of the Trp229 side chain shows distinct chemical shift perturbation in the ^1^H, ^15^N 2D HSQC spectrum during phosphorylation of the Puf3-IDR^1-258^ (Fig. 4D). Since this Trp lies within the DFW motif, we hypothesized that phosphorylation could alter Ccr4-Not binding and set out to experimentally test this by solution NMR. To compensate for increased heterogeneity in the spectra, we used the shorter Puf3-IDR^109-258^ construct for NMR analysis. Puf3-IDR^109-258^ is phosphorylated on the same residues as Puf3-IDR^1-258^ (except pS42, which is not present in this construct; Extended Data Fig. 10D-E), confirming the selectivity of the kinases. We titrated full-length Ccr4-Not into phosphorylated Puf3-IDR^109-258^ and observed less perturbation compared with unmodified Puf3-IDR^109-258^ (Extended Data Fig. 11A-B). Thus, phosphorylation likely reduces Ccr4-Not interaction with the Puf3 IDR.

To further understand the effect of phosphorylation on Puf3-IDR, we investigated the backbone dynamics of the IDR. We used NMR to analyze the rotational diffusion of Puf3-IDR by measuring its longitudinal (^15^N T1) and transverse (^15^N T2) relaxation times. Upon phosphorylation, the ^15^N T1/T2 ratio increases across the sequence, particularly around the DFW motif, indicating a decrease in local flexibility in the Ccr4-Not binding sites of the Puf3-IDR (Fig. 4E and Extended Data Fig. 11C). Thus, our data suggest that phosphorylation modulates both the charges and backbone dynamics of Puf3-IDR, leading to a reduced affinity for Ccr4-Not.

Next, we tested the effect of phosphorylation on the enzymatic activity of Ccr4-Not. We prepared differentially phosphorylated full-length Puf3 using three distinct phosphorylation timepoints (Fig. 4B) and assayed their ability to stimulate Ccr4-Not activity using *in vitro* deadenylation assays. Phosphorylation of Puf3 does not affect its RNA-binding capacity but it reduces its ability to stimulate Ccr4-Not dependent deadenylase activity in an additive manner (Fig. 4F and Extended Data Fig. 12). In summary, Puf3 is sequentially phosphorylated, which reduces its interaction with Ccr4-Not and its ability to stimulate deadenylation activity. This demonstrates that the *in vitro* enzymatic activity of Ccr4-Not towards specific RNAs can be tuned in a continuous manner. Since Puf3 is also regulated by phosphorylation *in vivo* ^25^, it likely tunes deadenylation in a rheostat-like manner in cells.

## Multivalency and phosphoregulation are common properties of Ccr4-Not binding IDRs

Most known RNA adaptors bind Ccr4-Not via IDRs ^1^ (Extended Data Fig. 13). We therefore tested other RNA adaptors for multivalent binding and phospho-regulation. For these experiments, we used a recombinant human CCR4-NOT complex ^27^ and human RNA adaptors.

Human PUM1 is a Pumilio RNA adaptor from the same family as *S. pombe* Puf3. An IDR in PUM1, as well as in Drosophila PUM, directly interacts with the CNOT1, CNOT2 and CNOT3 subunits of CCR4-NOT through a region that is proximal to the Pumilio domain ^15,16^. To test whether this IDR accounts for the stimulatory activity of human PUM1 in a fully reconstituted system, we purified N-terminal IDR deletions of PUM1 (Extended Data Fig. 14A). A truncation mutant containing a previously defined CCR4-NOT interacting region (PUM1^Δ587^) retained the ability to stimulate CCR4-NOT, but was ∼4-fold less efficient than the full-length protein (Fig. 5A and Extended Data Fig. 14B-C). Other truncations showed that the length of the IDR in the PUM1 construct correlates with the ability to stimulate deadenylation. This graded response is consistent with a multivalent binding mechanism involving interaction regions throughout the IDR, similar to yeast Puf3.

**Fig. 5.**
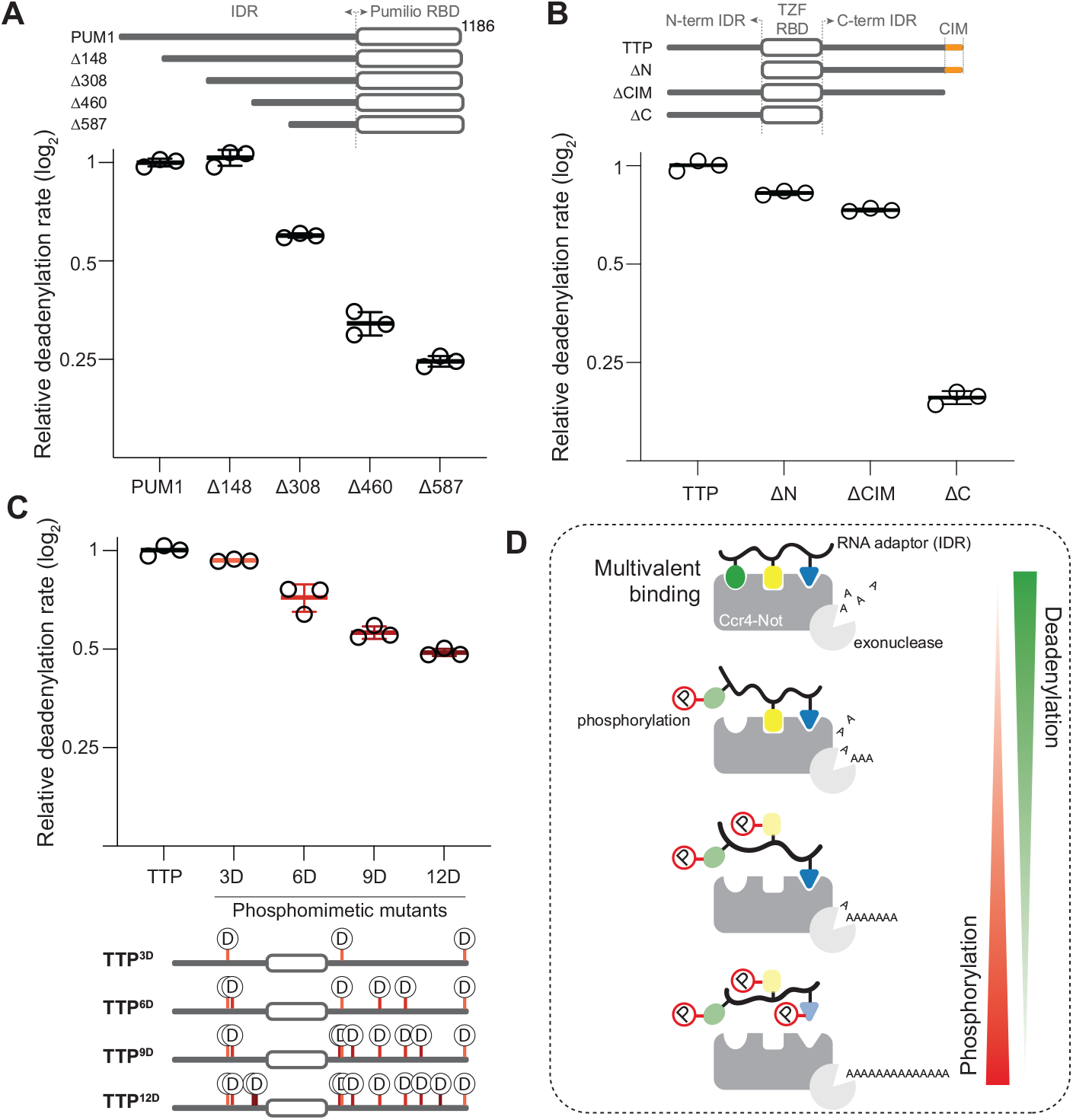
Multivalency can tune human CCR4-NOT mediated deadenylation with divergent RBPs. **(A)** Reconstitution of targeted deadenylation using recombinant full-length *H. sapiens* CCR4-NOT deadenylase complex and Pumilio homologue 1 (PUM1) constructs. Schematics of the PUM1 constructs are shown at the top. A 5’ fluorescently-labelled RNA containing an upstream PRE followed by a 30-nt poly(A) tail was used as a substrate. Reactions were started by addition of the deadenylase complex and resulting products were resolved on denaturing UREA PAGE gels. Experiments were performed in triplicate (open circles) and averaged (line), with error bars displaying the standard deviation. **(B)** Deadenylation assays performed as in panel A but in the presence of Tristetraprolin (TTP) constructs and with a substrate RNA containing an upstream AU-rich element (ARE). TTP contains both N- and C-terminal IDRs around a central tandem zinc-finger (TZF) RBD. A previously defined and structurally characterized CCR4-NOT interaction motif (CIM) is indicated in orange. (**C**) Deadenylation assays performed with TTP constructs containing the indicated number of phosphomimetic mutations. The locations of phosphomimetic mutations are indicated in the schematic diagrams below. (**D**) Model for multivalent interaction of RNA adaptors with CCR4-NOT, and regulation of phosphorylation. RNA-binding proteins tether Ccr4-Not to specific RNAs using a multivalent mode of binding through intrinsically disordered regions. Interactions can be tuned through phosphorylation across extended IDRs.

The human RNA adaptor protein TTP (also known as ZFP36) controls the stability of mRNAs involved in the inflammatory response ^28^. TTP interacts with CNOT1 through a conserved C-terminal SLiM ^12^ but also likely through additional regions in N- and C-terminal IDRs ^29^. TTP lacking the C-terminal SLiM shows a small but reproducible decrease in the ability to stimulate CCR4-NOT whereas TTP lacking the entire C-terminal IDR is much less active (Fig. 5B and Extended Data Fig. 15). Thus, additional elements in the IDR are important for maximal stimulation of deadenylation, presumably through binding to CCR4-NOT via a multivalent mechanism.

TTP is known to be regulated by phosphorylation during the inflammatory response but the exact roles of many of the phosphorylation sites are unknown ^30^. Given that these phosphorylation sites are distributed throughout the IDR we hypothesized that differentially phosphorylated TTP could exhibit similar behavior to Puf3 in tuning Ccr4-Not deadenylase activity. We purified full-length TTP variants with phosphomimetic mutations (Extended Data Fig. 16A) and tested their ability to stimulate the deadenylation activity of CCR4-NOT. Interestingly, CCR4-NOT exhibits a graded response to TTP phosphorylation, similar to Puf3 phosphorylation (Fig. 5C and Extended Data Fig. 16). Thus, multiple RNA adaptors likely bind Ccr4-Not through a multivalent interaction that can be regulated in a tunable manner to control mRNA deadenylation.

## Discussion

In this study, we demonstrate that RNA adaptor proteins including yeast Puf3, human PUM1 and human TTP interact with the Ccr4-Not deadenylase complex through a multivalent mechanism that can be regulated in a rheostat-like manner (Fig. 5D). Specifically, multiple regions within the IDRs of RNA adaptors have an additive effect on the ability to recruit Ccr4-Not to specific transcripts and to stimulate its deadenylation activity. The functional significance of such a mode of interaction lies in its regulatory potential: IDRs of RNA adaptors can be phosphorylated at specific sites, including within regions that bind to Ccr4-Not, thereby modulating their ability to regulate mRNA decay. In agreement with this, phosphorylation of Puf3 affects regulation of target transcripts in yeast ^25^ and phosphorylation of human TTP regulates mRNAs of the inflammatory response ^28^. Thus, in an updated tethering model, multivalent interaction and post-translational modification control the binding landscape between Ccr4-Not and specific transcripts. Given that regulation of poly(A) tail length is critical for mRNA half-life, phosphorylation can act as a rheostat to tune deadenylation rate and finely control RNA decay.

Previous studies had shown that there are several distinct binding pockets on Ccr4-Not that interact with SLiMs. However, these studies generally investigated short peptides in isolation and distinct subcomplexes of Ccr4-Not, without examination of the direct impact on deadenylation activity ^10,11,31,32^. Our strategy of using full-length RNA adaptors, the complete Ccr4-Not complex, and *in vitro* reconstitution, shows that individual SLiMs contribute to binding, but they are not sufficient for maximal stimulation of deadenylation. Instead, a combination of multiple regions of the IDR are required. The binding energy between each distinct SLiM and a Ccr4-Not binding pocket are likely small, consistent with a highly dynamic and shallow binding energy landscape ^33^. This could have the advantage of faster kinetics allowing greater accessibility to post-translational modification, a high capacity for regulation, and modularity, where multiple RNA adaptors work with a single enzymatic complex.

Many RNA adaptors interact with Not9, a very highly-conserved subunit of Ccr4-Not, suggesting that Not9 is a general IDR interaction hub. Since it interacts with many different RNA adaptors, Not9 has become evolutionary constrained. Other Ccr4-Not binding pockets (*e*.*g*. in CNOT1 and CNOT3) are also highly conserved. In contrast, IDRs normally have poor sequence conservation but retain function across evolutionarily distant proteins ^34,35^. Although the enzymatic machineries in gene expression (e.g. Ccr4-Not) are highly conserved, the sequence plasticity of IDRs allows evolutionary tuning of processes, in this case, regulation of mRNA decay. Interestingly, multivalency and IDRs are also important for interactions between the components of the decapping machinery ^36^. The interplay between different RNA adaptors, Ccr4-Not and the decapping machinery is currently not understood.

Our model for multivalent and rheostatic IDR interactions in mRNA decay has parallels with other processes in gene expression. For example, multivalency is essential for high affinity binding between transcription factor tandem activation domains (tADs) within Gcn4 and Mediator ^19^. In this case, individual hydrophobic SLiMs in Gcn4 have low affinity and specificity for Mediator. Multivalent binding increases the affinity and specificity of complex formation but there is not a single distinct ordered state – the tADs are highly dynamic, with multiple binding conformations and configurations. In addition, multi-site phosphorylation of p53 promotes binding to CREB-binding protein (CBP)/p300 ^18,37^ and decreases interaction with Mdm2 ^18^ in a graded manner. Thus, sequential phosphorylation of IDRs also regulates effector binding in other processes.

Overall, we demonstrate that RBPs mediate specificity by using a multivalent mechanism to interact with the Ccr4-Not complex that can be tuned by phosphorylation. Commonalities between deadenylation, decapping and transcription factors suggest that regulated multivalent interactions between IDRs in nucleic acid binding proteins and their effector complexes is likely a general strategy used to generate specificity within gene expression.

## Methods

### Molecular cloning and synthetic DNA construct design

All proteins described in this work were gene synthesized to optimize codon usage for *E. coli*. All PCR steps were performed using either Q5® or Phusion® hot start high fidelity polymerases and vectors were assembled isothermally using homology arms with NEBuilder® HiFi DNA Assembly Master Mix (NEB). Site-directed mutagenesis was performed by PCR using multiple overlapping fragments with primers incorporating substitutions before isothermal reassembly into vectors.

Gene synthesized subunits for both the fission yeast and human Ccr4-Not complexes cloned into pACEBac vectors with polyhedrin promoter based expression cassettes have been described previously ^27,38^. Cloning of *S. pombe* complexes was adapted to a modified BigBac system to streamline introduction of point mutants ^39,40^. Gene expression cassettes for Not1, Ccr4, Caf1 and Not9 in pACEBac1 vectors were PCR amplified and assembled into a single modified pBig1A vector. Cassettes for Not2, Not3 and Not4 in pACEBac1 were similarly assembled into a single modified pBig1B vector. pBig1 vectors were then assembled into a single modified pBig2AB vector using Pme1 digestion and isothermal assembly. Assembled complex vectors were then transformed into DH10 EMBacY cells for transposition into recombinant baculoviruses ^41^.

*S. pombe* Puf3 expression vectors have been previously described ^20^ and truncations were subcloned using specific PCR primers. *H. sapiens* PUM1 constructs were subcloned from a gene synthesized vector (GeneArt). Constructs for Tristetraprolin including phosphomimetic mutants were gene synthesized in expression vectors (Genscript). Kinase expression plasmids were subcloned using restriction digest from gene synthesized plasmids (Geneart).

### Recombinant protein expression and purification

#### Ccr4-Not complexes

Recombinant *S. pombe* and *H. sapiens* Ccr4-Not complex expression and purification has been previously described ^27,38^ but has been further optimized for this study. Isolated Bacmids were transfected into adherent Sf9 cells in 6-well plates using FuGene Transfect HD. Viral supernatant was harvested 72 h post-transfection and used immediately to infect expression cultures using a low titre strategy that is necessitated by the instability of the large complex viruses. 150 ml of Sf9 cells cultured in suspension in SF900-II SFM (Invitrogen) at 1.5-2 x 10^6^ cells/ml were infected with 1.5 ml of transfection supernatant. Cultures were then maintained at a cell density between 2-3 x 10^6^ by dilution with fresh medium until culture arrest ∼72 h post-infection. Cells were then cultured for a further 24-48 h before harvest by centrifugation at 1000 x rcf before pellets were gently washed with ice-cold PBS before flash freezing and storage at -80°C.

Frozen cell pellets were lysed by thawing in lysis buffer: 100 mM HEPES pH 8.0, 300 mM NaCl, 0.1% (v/v) Igepal CA-630, 2 mM DTT, protease inhibitor cocktail (Roche) and 0.4 mM PMSF. Lysate was cleared using ultracentrifugation at 200,000 *g* for 30 min and filtered using 0.65 μM PVDF membranes (Millipore). Clarified lysate was bound in batch to 1 ml Strep-Tactin Sepharose Resin (IBA Lifesciences) per litre of input culture for 1.5 h. Beads were washed with 50 mM HEPES (pH 8.0), 150 mM NaCl, 2 mM Mg(OAc)2, and 2 mM DTT before elution in buffer supplemented with 5 mM desthiobiotin (IBA Lifesciences). Eluate was directly loaded onto a HiTrap Q HP 5 ml column (Cytiva), and bound complexes eluted using a 12-column volume gradient. Peak fractions were collected, pooled and injected onto size exclusion chromatography using Superose 6 Prep Grade XK16/70 or 26/60 columns (Cytiva) equilibrated with 20 mM HEPES pH 8.0, 150 mM NaCl, 2 mM Mg(OAc)2, and 0.5 mM TCEP. Peak fractions were pooled and loaded onto a Resource Q 1 ml column and eluted using a 10 column volume gradient in 20 mM HEPES pH 8, ∼300 mM NaCl, 2 mM Mg(OAc)2, and 0.5 mM TCEP. This concentrated the complex to 2–5 mg/ml, which was then flash frozen in liquid nitrogen for storage at 80°C. For NMR experiments where complex at ∼10 mg/ml was required, complex was supplemented with 10% (w/v) glycerol before further concentration using Amicon Ultra-4 50K MWCO centrifugal filter units.

Complex concentration was determined using UV-visible spectrophotometry. Stoichiometry and purity were determined at each purification step using NuPAGE 4-12% gradient SDS-PAGE gel analysis. Complex stability and dispersity were monitored using light scattering, mass photometry and differential scanning fluorimetry.

#### Kinases

Recombinant Sck1 and Ksg1 carrying N-terminal twin-SII tags were overexpressed in Sf9 cells from baculoviruses transposed from pACEBac1 vectors. Bacmids were prepared and transfected into adherent Sf9 cells as for Ccr4-Not complexes and supernatant containing P1 virus harvested at 72 h post-transfection. Virus was then amplified in suspension culture by infecting cells at 1.5 x 10^6^ cells/ml density with 1:100 (v/v) P1 virus. Cell density was maintained by dilution with fresh media until cultures arrested, and viral P2 supernatant collected at 72 h post-infection. P2 virus was used to infect suspension expression cultures at 2.5 x 10^6^ cells/ml. Cells expressing Ksg1 were harvested at 48 h post-infection because of protein toxicity and Sck1 harvested 60 h post-infection using centrifugation as detailed for Ccr4-Not complexes.

Frozen cell pellets were lysed by thawing in detergent as detailed for Ccr4-Not complexes. Lysate was clarified and kinases purified using affinity purification as described for complexes above. Eluates were then directly injected onto a HiTrap Q HP 5 ml column (Cytiva), and bound protein eluted using a 15-column volume gradient. Peak fractions were concentrated using Amicon Ultra-15 30K MWCO centrifugal filter units.

#### Pumilio protein constructs

All *S. pombe* Puf3 and *H. sapiens* constructs for assays were overexpressed as MBP fusion proteins from a modified pMAL-c5x vector where the thrombin cleavage site was replaced with a 3C protease site. NMR constructs were expressed from modified pMAL-c5x vector carrying 3C-cleavable N-terminal MBP and C-terminal lipoyl tags. Puf3 protein constructs for binding assays were expressed from a modified pGEX vector carrying N-terminal GST and C-terminal lipoyl tags. Vectors was transformed into in-house prepared chemically competent BL21star (DE3) cells and positive colonies were directly used to inoculate expression cultures in 2xTY medium or minimal medium for isotopic labelling. MBP tagged proteins for assays were expressed in 2xTY medium, grown to OD600nm 0.8-1 and induced with 0.5 mM IPTG for 4 h at 25°C. GST tagged fragments were induced overnight at 18°C. Uniformly labelled proteins were expressed in M9 minimal media (6 g/l Na2HPO4, 3 g/l KH2PO4, 0.5 g/l NaCl) supplemented with 1.7 g/l yeast nitrogen base without NH4Cl and amino acids (Sigma Y1251). 1 g/l ^15^NH4Cl and 4 g/l unlabeled glucose were supplemented for ^15^N labelling. Unlabeled glucose was replaced with 3 g/l ^13^C-glucose for ^13^C/^15^N double-labelled samples. Cells were harvested by centrifugation and stored at -80°C before lysis.

Cell pellets were thawed in 5 volumes of lysis buffer containing 50 mM HEPES pH 8, 300 mM NaCl, 0.5 mM TCEP, 2 mM EDTA, 0.5 mM PMSF, Protease inhibitor cocktail (Roche), 10 μM Leupeptin and 2 μM Pepstatin. Cells were lysed by sonication in 50 ml batches for 30 sec total time with 30% amplitude using a 5 mm microtip on a VCX750 system (Sonics). Lysates were cleared by ultracentrifugation at 150,000*g* for 30 min before filtration through 0.65 μM filters. MBP tagged protein was subsequently bound in batch for 1 h to 1 ml of Amylose resin (NEB) per litre of input culture (typically 4 ml total resin). The resin lysate suspension was then transferred to a 2.5×20 econo-column (Biorad) and resin collected before the bed was washed with 3 x 100 ml of wash buffer (50 mM HEPES pH 8, 300 mM NaCl, 0.5 mM TCEP, 2 mM EDTA and 5 μM Leupeptin). Protein was then eluted using elution buffer (20 mM PIPES pH 6.8, 200 mM NaCl, 50 mM Maltose, 0.01% (w/v) Brij-35 and 0.5 mM TCEP).

Eluted proteins were immediately loaded onto a 5 ml HiTrap Heparin HP column equilibrated in 20 mM HEPES pH 6.8, 100 mM NaCl and 0.5 mM TCEP and bound protein was eluted using a 15-column volume linear gradient. Peak fractions were pooled and injected onto a Superdex 200 26/60 prep grade size exclusion column and proteins eluted isocratically in 20 mM HEPES pH 7.5, 150 mM NaCl and 0.5 mM TCEP. For *S. pombe* Puf3 constructs, peak fractions were pooled and concentrated to ∼5 mg/ml using Amicon Ultra-15 50K MWCO centrifugal filter units before flash freezing for storage at -80°C. For *H. sapiens* PUM1 constructs, pooled fractions from size exclusion were loaded onto a 1 ml RESOURCE S column equilibrated in 20 mM Bis-Tris pH 6, 50 mM NaCl, 0.5 mM TCEP and 10% (w/v) glycerol. Protein was then eluted using a 10 CV linear gradient and peak fractions flash frozen for storage at -80°C.

#### Tristetraprolin protein constructs

All *H. sapiens* Tristetraprolin (TTP) constructs for assays were overexpressed as MBP fusion proteins from a modified pMAL-c5x vector where the thrombin cleavage site was replaced with a 3C protease site. Proteins were expressed in 2xTY medium and supplemented with 10 μM final concentration ZnSO4 on induction with 0.5 mM IPTG for 4 h at 25°C. TTP constructs were harvested and lysate prepared as for Pumilio proteins. Proteins were bound in batch to Amylose resin and washed with 3 x 100 ml wash buffer (50 mM HEPES pH 8, 300 mM NaCl, 10 μM ZnAc, 0.5 mM TCEP, 2 mM EDTA and 5 μM Leupeptin). Bound proteins were then eluted with 20 mM PIPES pH 7, 200 mM NaCl, 0.5 mM TCEP, 1 μM ZnAc, 0.1% Brij 35. Downstream purification steps of cation exchange and size exclusion chromatography were then performed as for Pumilio proteins above.

### *In vitro* Protein Phosphorylation

Purified Sck1, along with its activation loop kinase Pdk1, were produced as described above. Phosphorylated Puf3-IDR^1-258^ and Puf3-IDR^109-258^ for NMR analysis was produced using a dialysis method: Puf3 IDR constructs were purified according to the relevant section above up until the first HiTrap Heparin HP column step. Peak fractions were pooled and mixed with a kinase mastermix in a 5:1 molar ratio. Kinase mastermix contained equimolar amounts of Sck1 and Pdk1 and was preincubated for 15 min with 5 mM ATP and 20 mM MgCl2 before mixing with Puf3. 10 ml phosphorylation reactions were dialyzed 50-fold using Slide-A-Lyzer 10K MWCO G2 cassettes against 500 ml of phosphorylation buffer: 20 mM HEPES pH 7.4, 10 mM NaCl, 0.5 mM TCEP, 5 mM ATP (99% Roche) and 20 mM MgCl2. Phosphorylation was typically allowed to proceed under dialysis for 18 h at 4°C. Reaction mixture was then applied to a HiTrap Heparin HP 5ml column to remove kinase and unphosphorylated IDR. Flow through was collected and applied to a HiTrap Q HP 5 ml column and phosphorylated species eluted using a 15 column volume gradient. Peak fractions were then injected onto a HiLoad Superdex 200 26/60 prep grade column equilibrated in 20 mM HEPES pH 8, 100 mM NaCl and 0.5 mM TCEP. Phosphorylated IDRs were subsequently resolved on a Capto HisRes Q 5/50 using a 50 column volume gradient before peak fractions were immediately dialyzed into NMR buffer (see NMR methods).

Phosphorylation time course experiments with full-length MBP-Puf3 were performed in 0.5 ml volumes in 10K MWC Slide-A-Lyzer™ MINI dialysis devices in phosphorylation buffer: 20 mM HEPES pH 7.4, 50 mM NaCl, 4 mM ATP, 20 mM MgCl2, 0.5 mM TCEP. Puf3 concentration was adjusted to 32.5 μM and mixed with 6 μM final concentration kinase mixture to start reactions. Phosphorylation time courses were performed at room temperature under dialysis with rapid stirring against a 50-fold excess volume of phosphorylation buffer. 100 μl aliquots at desired time points were withdrawn and immediately desalted into 20 mM HEPES pH 8, 100 mM KCl, 0.5 mM EDTA and 0.5 mM TCEP using PD SpinTrap G-25 spin column with a 25 μl stacker volume. Phosphorylated proteins were then immediately used in deadenylation activity assays, for mass spectrometry or RNA-binding assays.

### Deadenylation assays

Assays using *S. pombe* Ccr4-Not were performed as previously described with some modifications ^20,38^. Substrate RNA was synthesized with a 5’ 6-FAM fluorophore and contained a Pumilio response element (PRE) embedded within a 20-nt non-poly(A) region, upstream of a 30-nt poly(A) tail. Recombinant Ccr4-Not complexes were diluted to either 0.5 μM (10x stock) for targeted assays (in the presence of an RNA adaptor) or 1 μM for assays alone (no RNA adaptor) in 20 mM HEPES pH 8, 300 mM NaCl, 2 mM Mg Acetate and 0.5 mM TCEP. Puf3 constructs were diluted to 2.5 μM (10x stock) in 20 mM HEPES pH 8, 150 mM KCl and 0.5 mM TCEP. Assays were performed in 100 μl total volume in a thermally controlled block at 22°C. RNA (final concentration 200 nM) was preincubated with 250 nM Puf3 construct for 15 min in deadenylation buffer (20 mM PIPES pH 6.8, 10 mM KCl, 2 mM Mg acetate and 0.1 mM TCEP). Reactions were then started by the addition of 10x Ccr4-Not stock to a final concentration of either 50 nM or 100 nM (for control reactions in Extended Data Fig. 8). 7.5 μl samples at indicated time points were withdrawn and mixed with an equivalent volume of loading dye (95% Formamide, 4 mM EDTA, 0.01 w/v bromophenol blue). Deadenylated species were resolved on denaturing TBE (Tris-borate-EDTA) 20% polyacrylamide (19:1) gels containing 7 M urea and run at 400 V in 1 × TBE running buffer for 30 min. Gels were visualized using an Amersham Typhoon 5 system (Cytiva) using 488 nm excitation laser and 525 nm 20 nm bandpass emission filter.

Assays using *H. sapiens* CCR4-NOT were performed as previously described but with conditions optimized for the human complex. 10x CCR4-NOT complex and 10x RNA-binding protein stocks were prepared at 0.5 μM and 2.5 μM respectively in the same buffers as for the *S. pombe* complex above. Assays with TTP were performed with an RNA containing an upstream ARE element instead of the PRE element as detailed above, also containing a 30-nt poly(A) tail and an 5’ 6-FAM fluorophore for visualization. RNA at a final concentration of 200 nM was preincubated with indicated 250 nM RNA-binding protein construct in deadenylation buffer optimized for the human complex (20 mM HEPES pH 7.8, 80 mM NaCl, 2 mM MgAc and 0.1 mM TCEP final concentration). Reactions were started with the addition of 10x CCR4-NOT complex and samples withdrawn, reaction stopped and resolved on denaturing PAGE gels as with *S. pombe* complex assays described above.

Assays were cropped and contrast adjusted linearly in Photoshop (Adobe) before being imported into Image J. Complete lane profiles were integrated and intensity values obtained for both total substrate and deadenylated substrate per lane/time point. Relative deadenylated fractions were calculated, plotted in Prism (GraphPad) as a function of time and fitted with a logistic function to yield fitted densitometry curves. Poly(A) tail half-lives were used to calculate relative reactivities compared with stimulation of deadenylation by the full-length wild-type RBP. Assays and quantification were performed 3 times for every construct.

### Fluorescence anisotropy

Binding data for *S. pombe* Puf3 constructs to Ccr4-Not was obtained using the complex as a fluorescent probe. Ccr4-Not was site specifically FAM-labelled on the N-terminus of the Not9 subunit using a YbbR tag ^42^. CoA-Fluorescein conjugates were synthesized from CoA and fluorescein maleimide before RP-HPLC purification, lyophilization and LC-MS validation. YbbR-tagged Ccr4-Not complexes were prepared according to the molecular cloning and purification sections above. An in-frame YbbR tag was introduced at the N-terminus of Not9 through PCR and assembled into a vector containing the full Ccr4-Not complex for baculovirus mediated overexpression. Complex was purified and labelled by incubation of Ccr4-Not with 5-fold molar excess of CoA-fluorescein and 1:100 Sfp enzyme (prepared in-house) in 20 mM HEPES pH 8, 150 mM NaCl, 0.5 mM TCEP and 5 mM MgCl2. Excess dye and Sfp was then removed using size-exclusion chromatography and the complex was further purified by anion exchange chromatography as detailed in the complex purification section above. Binding was measured by titration of increasing concentrations of Puf3 against 25 nM final concentration labelled Ccr4-Not in binding buffer: 20 mM HEPES pH 8, 100 mM NaCl and 0.5 mM TCEP, in a total volume of 20 μl using 384-well black bottom round bottomed low-volume non-binding surface assay microplates (Corning). Fluorescence anisotropy was measured using a PHERAstar Plus (BMG Labtech). Binding curves were fitted using a quadratic function, taking ligand depletion into account, to calculate dissociation constants.

For RNA-binding, fluorescently labeled RNA oligonucleotides were used, containing a PRE element embedded within a 20-nt RNA. Two-fold protein dilution series were prepared at 10 × concentration in dilution buffer (20 mM HEPES pH 7.5, 100 mm NaCl, 0.5 mm TCEP). Protein was incubated at room temperature for 30 min with 0.1 nm 5’ FAM-labelled RNA (synthesized by IDT) in buffer containing 20 mM HEPES pH 7.5, 150 mM NaCl, 0.1 mM TCEP, in a total volume of 100 μl using 384-well low-flange black flat bottom nonbinding surface microplate (Corning). Fluorescence anisotropy was measured using a PHERAstar Plus (BMG Labtech). Dissociation constants were estimated using a quadratic function Prism 6.0 (GraphPad software). Error bars indicate the standard deviation of 2 technical replicates.

### Crosslinking mass spectrometry

Sulfo-NHS-Diazirine (Sulfo-SDA) based UV-induced crosslinking was performed both on *S. pombe* Ccr4-Not complex alone and with 1.5-fold molar excess Puf3. Sulfo-SDA stocks were freshly dissolved in ultrapure water at 100 mM before being added to a final concentration of 0.5, 0.1 and 0.05 mM in three separate crosslinking reactions with a total of 100 μg of complex adjusted to 0.2 μg/μl. Reactions were incubated on ice for 30 min before the diazirine moiety was activated by irradiation under a 365 nm UV lamp for 20 min. Bands corresponding to crosslinked complex in the 0.5 mM samples were excised from SDS-PAGE gels. Lower concentration (0.1 and 0.05 mM) samples were then mixed, precipitated using ice cold acetone and proteins pelleted and dried.

The precipitated protein samples were resolubilized in digestion buffer (8 M urea in 100 mM ammonium bicarbonate) to an estimated protein concentration of 1 mg/ml. Disulfide bonds in the samples were reduced by adding dithiothreitol (DTT) to a final concentration of 5 mM, followed by incubation at room temperature for 30 min. The free sulfhydryl groups were then alkylated by adding iodoacetamide to a final concentration of 15 mM and incubating at room temperature for 20 min in the dark. After alkylation, the excess iodoacetamide was quenched by adding DTT to a final concentration of 10 mM.

Next, the protein samples were digested with LysC at a 50:1 (w/w) protein-to-protease ratio at room temperature for four hours. The samples were then diluted with 100 mM ammonium bicarbonate to reduce the urea concentration to 1.5 M. Trypsin was added at a 50:1 (w/w) protein-to-protease ratio to further digest the proteins for 15 hours at room temperature. The resulting peptides were desalted using C18 StageTips ^43^.

For each sample, the resulting peptides were fractionated using size exclusion chromatography (SEC) to enrich for crosslinked peptides ^44^. Peptides were separated using a Superdex™ 30 Increase 3.2/300 column (GE Healthcare) at a flow rate of 10 μl/min. The mobile phase consisted of 30% (v/v) acetonitrile and 0.1% trifluoroacetic acid. The first seven peptide-containing fractions (50 μl each) were collected, and the first two fractions were combined. The solvent was removed using a vacuum concentrator, and the fractions were then analyzed by LC-MS/MS.

LC-MS/MS analysis was performed using an Orbitrap Fusion Lumos Tribrid mass spectrometer (Thermo Fisher Scientific) connected to an Ultimate 3000 RSLCnano system (Dionex, Thermo Fisher Scientific). Each SEC fraction was resuspended in 1.6% (v/v) acetonitrile and 0.1% (v/v) formic acid and analyzed by LC-MS/MS. Each SEC fraction was analyzed in duplicate by LC-MS/MS. Peptides were injected onto a 50-cm EASY-Spray C18 LC column (Thermo Scientific), operated at 50 °C. Mobile phase A consisted of water with 0.1% (v/v) formic acid, and mobile phase B consisted of 80% (v/v) acetonitrile with 0.1% (v/v) formic acid. Peptides were loaded and separated at a flow rate of 0.3 μl/min. Separation was achieved using linear gradients, increasing from 2% to 55% B over 92.5 min. The gradient slope was optimized for each SEC fraction. Following separation, the B content was ramped to 95% over 2.5 min. Eluted peptides were ionized by an EASY-Spray source (Thermo Scientific) and introduced directly into the mass spectrometer.

The MS data were acquired in data-dependent mode. In each 2.5-sec acquisition cycle, the full scan mass spectrum was recorded in the Orbitrap at a resolution of 120,000. Ions with a charge state of 3+ to 7+ were isolated and fragmented using higher-energy collisional dissociation (HCD). For each isolated precursor, a collision energy of 26%, 28%, or 30% was applied, depending on the m/z and charge state of the precursor ^45^. The fragmentation spectra were then recorded in the Orbitrap at a resolution of 60,000. Dynamic exclusion was enabled with a single repeat count and a 30-sec exclusion duration.

For MS raw data processing, MS2 peak lists were generated using the MSConvert module in ProteoWizard (version 3.0.11729). Precursor and fragment m/z values were recalibrated. Crosslinked peptides were identified using the xiSEARCH software (version 2.0) (https://www.rappsilberlab.org/software/xisearch) ^46^. The data from the apo Ccr4-Not complex and Ccr4-Not+Puf3 samples were analyzed separately. Peak lists were searched against the protein sequences of the subunits of the *Schizosaccharomyces pombe* Ccr4-Not complex, and for the Ccr4-Not+Puf3 sample, the sequence of Puf3 was included. The reversed protein sequences were used as decoys for error estimation.

The following parameters were applied during the search: MS accuracy at 3 ppm; MS2 accuracy at 5 ppm; enzyme specificity set to trypsin with full tryptic cleavage; up to three missed cleavages allowed; and allowance for up to two missing monoisotopic peaks. Carbamidomethylation on cysteine was set as a fixed modification, while oxidation on methionine was set as a variable modification. SDA was specified as the crosslinker, with reaction specificity for lysine, serine, threonine, tyrosine, and protein N-termini on the NHS ester end, and any amino acid for the diazirine end. SDA loop links and hydrolyzed SDA on the diazirine end were also considered as variable modifications.

Crosslinked peptide candidates from each sample were filtered using xiFDR (version 2.2.betaB) ^47,48^ with a requirement of a minimum of three matched fragment ions per crosslinked peptide, including at least two ions containing a crosslinked residue. A false discovery rate of 1% at the residue-pair level was applied, with the ‘boost between’ option enabled. The mass spectrometry proteomics data have been deposited to the ProteomeXchange Consortium via the PRIDE ^49^ partner repository with the dataset identifier PXD055147 and 10.6019/PXD055147.

### Intact mass spectrometry

Proteins were desalted and buffer exchanged into ammonium acetate (50 mM) and formic acid (0.1 % v/v) using Microcon molecular weight cut-off filters (Millipore, US). Denatured proteins were washed 5 times before being diluted to a concentration of approximately 2 μg/μl. Proteins were analyzed via direct infusion intact mass spectrometry. All analysis was carried out on a Synapt G2-Si (Waters, UK) using a pulled borosilicate glass emitter coupled to a nano-electrospray (n-ESI) ion source in positive ion mode. The data were acquired using 5 second scan rates and the spectra were processed using UniDec ^50^ deconvolution.

### NMR Spectroscopy

All NMR samples were dialyzed against 2 L of the following buffers before data collection: For backbone assignment experiments, the samples were prepared in 20 mM PIPES pH 6.8, 50 mM NaCl, 2 mM TCEP and 0.02% sodium azide. For Ccr4-Not binding experiments, the samples were dialyzed into 20 mM PIPES pH 6.8, 250 mM NaCl, 2 mM TCEP and 0.02% sodium azide. For phosphorylation time courses, the samples were prepared in 20 mM PIPES pH 6.8, 50 mM NaCl, 2 mM TCEP, 0.02% sodium azide, 1 mM ATP and 2 mM MgCl2. Time courses were started with the addition of 1:5 Kinase. 5 % D2O was supplemented to all NMR samples.

Most experiments were performed using an in-house Bruker 800MHz Avance III spectrometer, equipped with a triple resonance TCI cryoProbe. Access to a Bruker 950 MHz Avance III spectrometer located at the MRC Biomedical NMR Centre (Francis Crick Institute) provided experiments with increased resolution and sensitivity. They were especially beneficial for the unambiguous assignment of exchange broadened residues. Unless otherwise specified, spectra were collected using a 75 μM sample in 5 mm tube at 278 K.

Backbone resonances of all Puf3-IDR constructs (Puf3-IDR^1-258^, Puf3-IDR^109-258^ and phosphorylated Puf3-IDR^109-258^) were obtained with BEST-TROSY versions of triple resonance experiments: HNCO, HN(CA)CO, HNCACB and HN(CO)CACB (Bruker). All 3D datasets were collected with non-uniform sampling at 10-25% and processed in MddNMR using compressed sensing reconstruction ^51^. To obtain a complete assignment of the backbone resonances of prolines and residues that underwent rapid solvent exchange, we assigned additional non-amide resonances using ^1^H start versions of ^13^C-detected CON, CACON and CANCO (Bruker). Backbone resonances were assigned using in-house scripts and Mars ^52^. Topspin 4.1.1 (Bruker) was used for processing and NMRFAM-Sparky 1.47 was used for data analysis ^53^.

For binding studies with Ccr4-Not, the relative peak intensities were normalized to the C-terminal residue of the construct (Gln from cleavage scar) and expressed as PIbound/PIfree, with PIfree and PIbound being the peak heights of the free and bound forms, respectively.

^15^N longitudinal (T1) and transverse (T2) relaxation times were measured using INEPT based pseudo-3D experiments with an initial recovery delay of 5s. Temperature compensation delays during the T2 experiments were used to account for sample heating effects. Mixing times used for T1 measurements were 0.01, 0.02, 0.04, 0.08, 0.12, 0.16, 0.32, 0.64, 1.28, 1.60, 2.00 and 0.01 s. Mixing times used for T2 measurements were 8.48, 16.96, 33.92, 50.88, 67.84, 101.76, 135.68, 169.6, 203.52, 237.44, 271.36 and 8.48 ms. Peak height analysis was performed in NMRFAM-Sparky 1.47 ^53^.

## Acknowledgements

We thank Pablo Alcón (MRC-LMB) for advice on protein labelling; J. G. Shi (MRC-LMB) for support with baculovirus; Meng Su for help with RP-HPLC; and all members of the Passmore group for useful discussions and advice. This work was supported by the MRC as part of UK Research and Innovation, MRC file reference number MC_U105192715 (L.A.P.); the European Union’s Horizon 2020 research and innovation program (ERC Consolidator grant agreement 725685 [to L.A.P.]); the Deutsche Forschungsgemeinschaft (DFG, German Research Foundation) under Germany’s Excellence Strategy – EXC 2008 – 390540038 – UniSysCat; and the Wellcome Trust through a Discovery Award (227434) and core funding for the Wellcome Centre for Cell Biology (203149). NMR was supported by the Francis Crick Institute through provision of access to the MRC Biomedical NMR Centre. The Francis Crick Institute receives its core funding from Cancer Research UK (CC1078), the UK Medical Research Council (CC1078), and the Wellcome Trust (CC1078).

## Author contributions

J.A.W.S. and L.A.P. conceived the study and designed the experiments. J.A.W.S. purified proteins, performed *in vitro* assays, and G.L. and S.A. assisted. C.W.H.Y., J.A.W.S. and S.M.V.F. performed NMR. Z.A.C., T.M., L.S., F.J.O’R. and J.R. performed mass spectrometry experiments. J.A.W.S. and L.A.P. wrote the manuscript and prepared figures with input from all authors. L.A.P. supervised the project.

## Competing interests

The authors declare that they have no competing interests.

## Notes

### Competing Interest Statement

The authors have declared no competing interest.

