## Supplementary Figures for "Phosphorylation-dependent tuning of mRNA deadenylation rates"

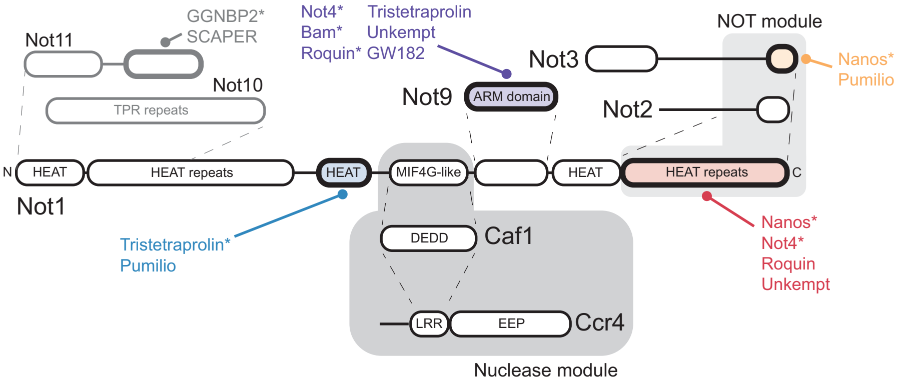


**Extended Data Fig. 1: Architecture of Ccr4-Not interactions with RNA adapters.** Schematic diagram showing known interactions between domains of Ccr4-Not and RNA-binding protein short linear motifs (SLiMs). Structured domains are depicted as rounded rectangles and linkers or disordered regions are shown as connecting lines. All core subunits of the complex assemble around the Not1 scaffold protein (interacting regions indicated with dashed lines) except Ccr4, which is bridged via Caf1. Many subunit interactions have been studied and some interfaces have been described through crystal structures (asterisks). Colored domains contain known interaction surfaces with SLiMs.

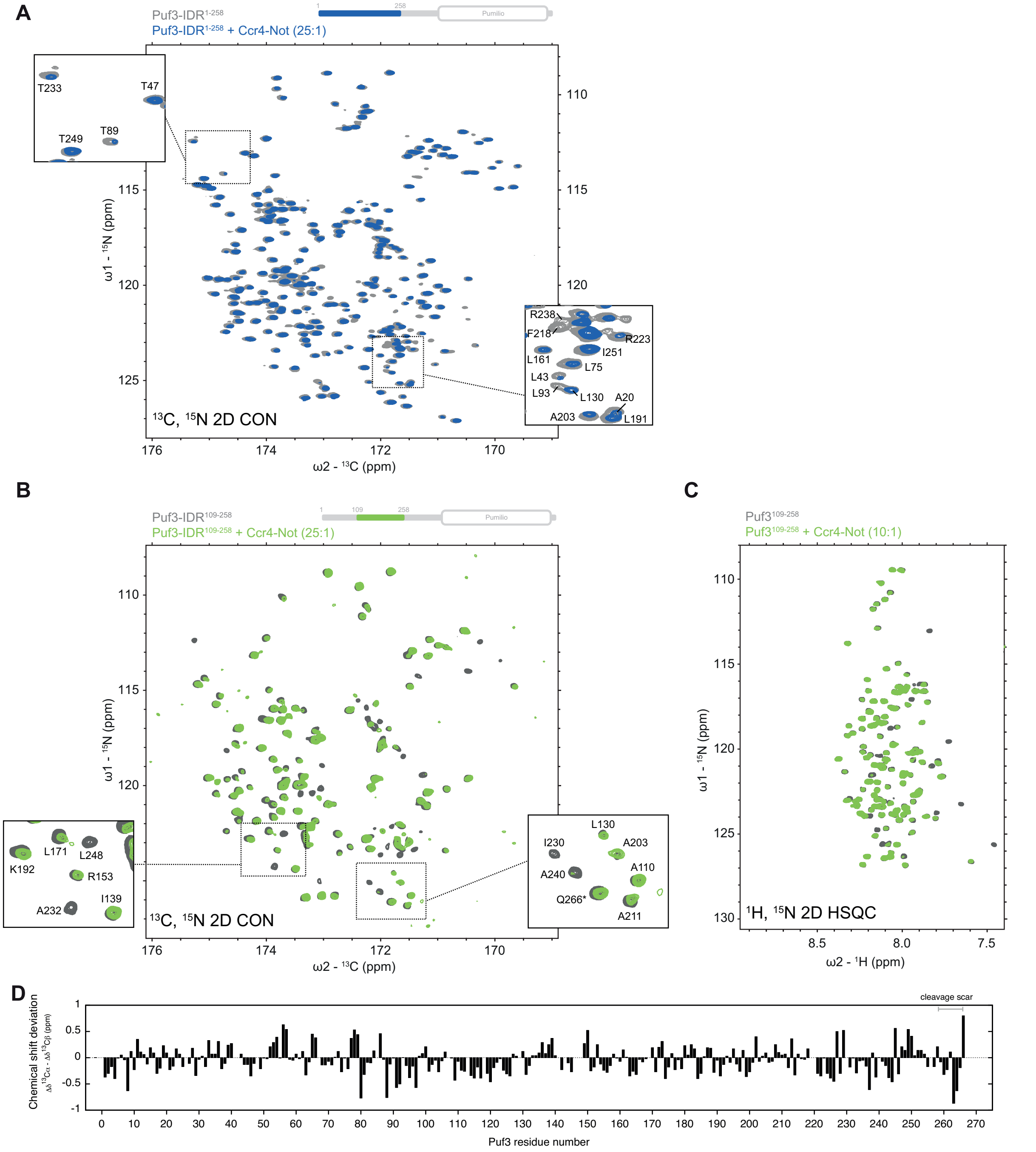


**Extended Data Fig. 2: NMR analysis of Puf3 IDR constructs with Ccr4-Not.** Spectra for Puf3-IDR$^{1-258}$ and Puf3-IDR$^{109-258}$ were recorded in the presence of either 1:10 or 1:25 full-length recombinant Ccr4-Not complex. (**A**) $^{13}$C 2D CON spectrum overlay of 75 μM Puf3-IDR$^{1-258}$ with (blue) or without (grey) 3 μM unlabeled Ccr4-Not. (**B**) $^{13}$C 2D CON spectra of 75 μM Puf3-IDR$^{109-258}$ with (green) or without (grey) 3 μM unlabeled Ccr4-Not. (**C**) $^{1}$H,$^{15}$N 2D-HSQC of 75 μM Puf3-IDR$^{109-258}$ with (green) or without (grey) 7.5 μM unlabeled Ccr4-Not. (**D**) Secondary chemical shift analysis (Δδ13Cα - Δδ13Cβ) for Puf3-IDR$^{1-258}$. Positive values would suggest residual helicity within that region. These data show that Puf3-IDR$^{1-258}$ lacks substantial secondary structure.

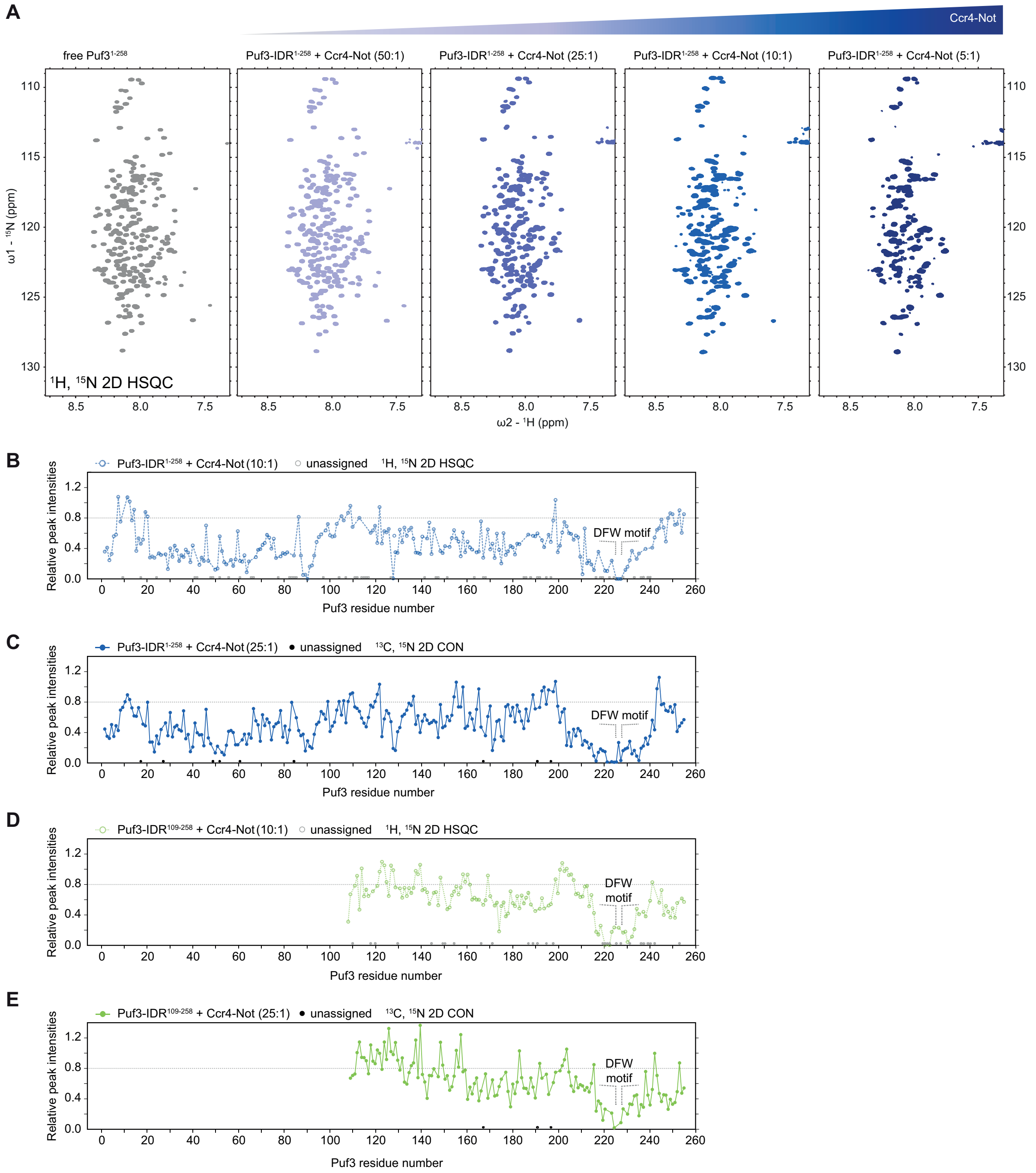


**Extended Data Fig. 3: Mapping the interaction of Ccr4-Not on Puf3.** (**A**) $^{1}$H,$^{15}$N 2D-HSQC of 75 µM Puf3-IDR$^{1-258}$ in the presence of the indicated concentrations of unlabeled Ccr4-Not. (**B**) Relative peak intensities of Puf3-IDR$^{1-258}$ in the presence of 1:10 Ccr4-Not (compared to Puf3-IDR$^{1-258}$ alone) from $^{1}$H,$^{15}$N 2D-HSQC calculated from spectra in Fig. 1C. (**C**) Relative peak intensities of Puf3-IDR$^{1-258}$ in the presence of 1:25 Ccr4-Not (compared to Puf3-IDR$^{1-258}$ alone) calculated from $^{13}$C 2D CON spectra in Extended Data Fig. 2A. Panels B and C are reproduced from Fig. 1D. (**D**) Relative peak intensities of Puf3-IDR$^{109-258}$ in the presence of 1:10 Ccr4-Not (compared to Puf3-IDR$^{109-258}$ alone) calculated from $^{1}$H,$^{15}$N 2D-HSQC spectra in Extended Data Fig. 2C. (**E**) Relative peak intensities Puf3-IDR$^{109-258}$ in the presence of 1:25 Ccr4-Not (compared to Puf3-IDR$^{109-258}$ alone) from $^{13}$C 2D CON plotted from spectra in Extended Data Fig. 2B.

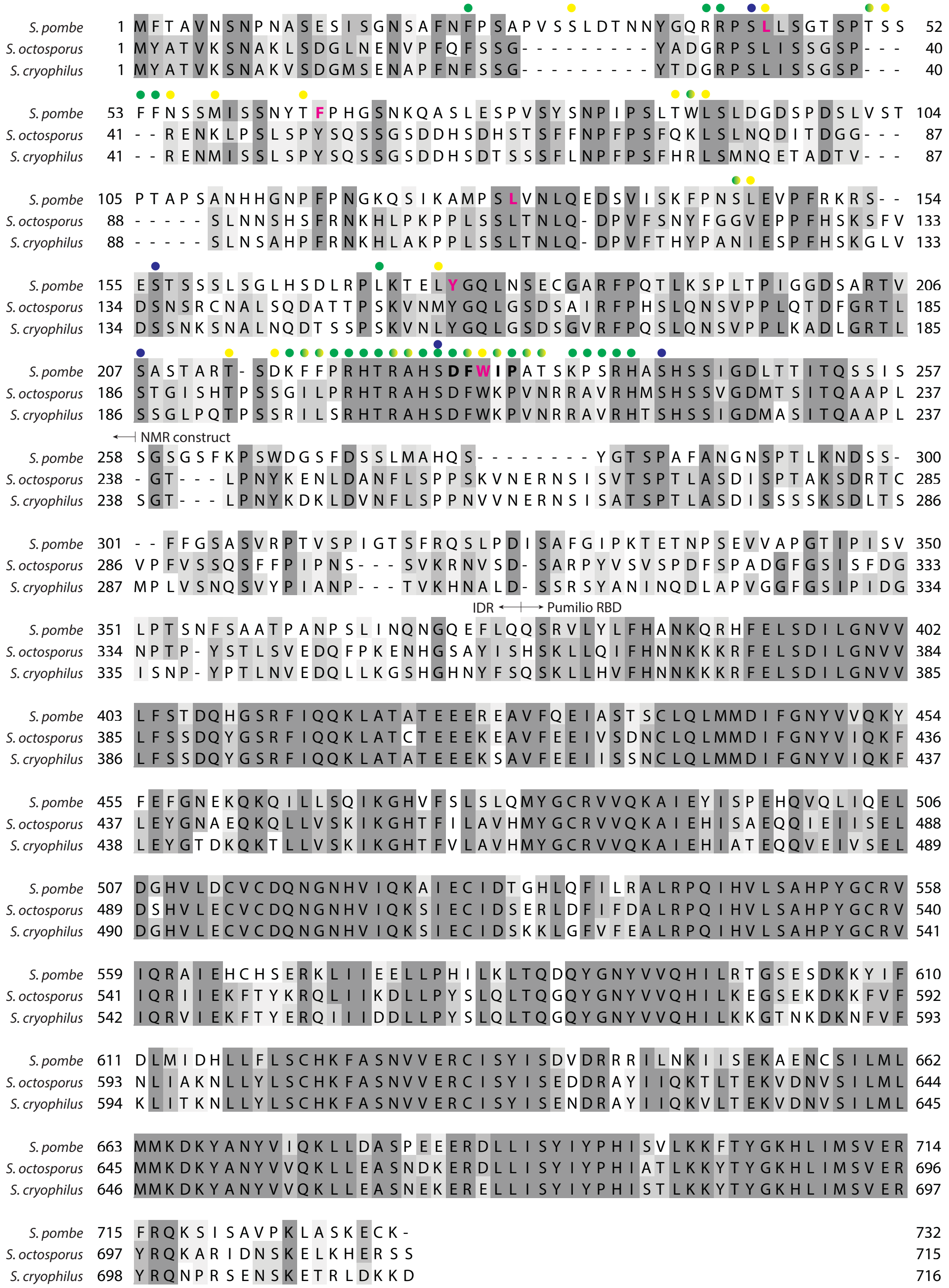


**Extended Data Fig. 4: Sequence alignments of Puf3.** Puf3 protein homologues within the *Schizosaccharomyces* genus. IDR and Pumilio RNA binding domain (RBD) boundaries are indicated. Residues that are exchange broadened in the presence of Ccr4-Not (<10% intensity relative to IDR alone) are shown with yellow dots for $^{1}H$,$^{15}N$ 2D-HSQC, green dots for $^{13}C$ 2D CON, or green/yellow dots for both. Residues in the DFW motif are in bold (black) and residues mutated in Fig. 2 are in bold magenta. Phosphorylation sites characterized in this study are indicated with blue dots.

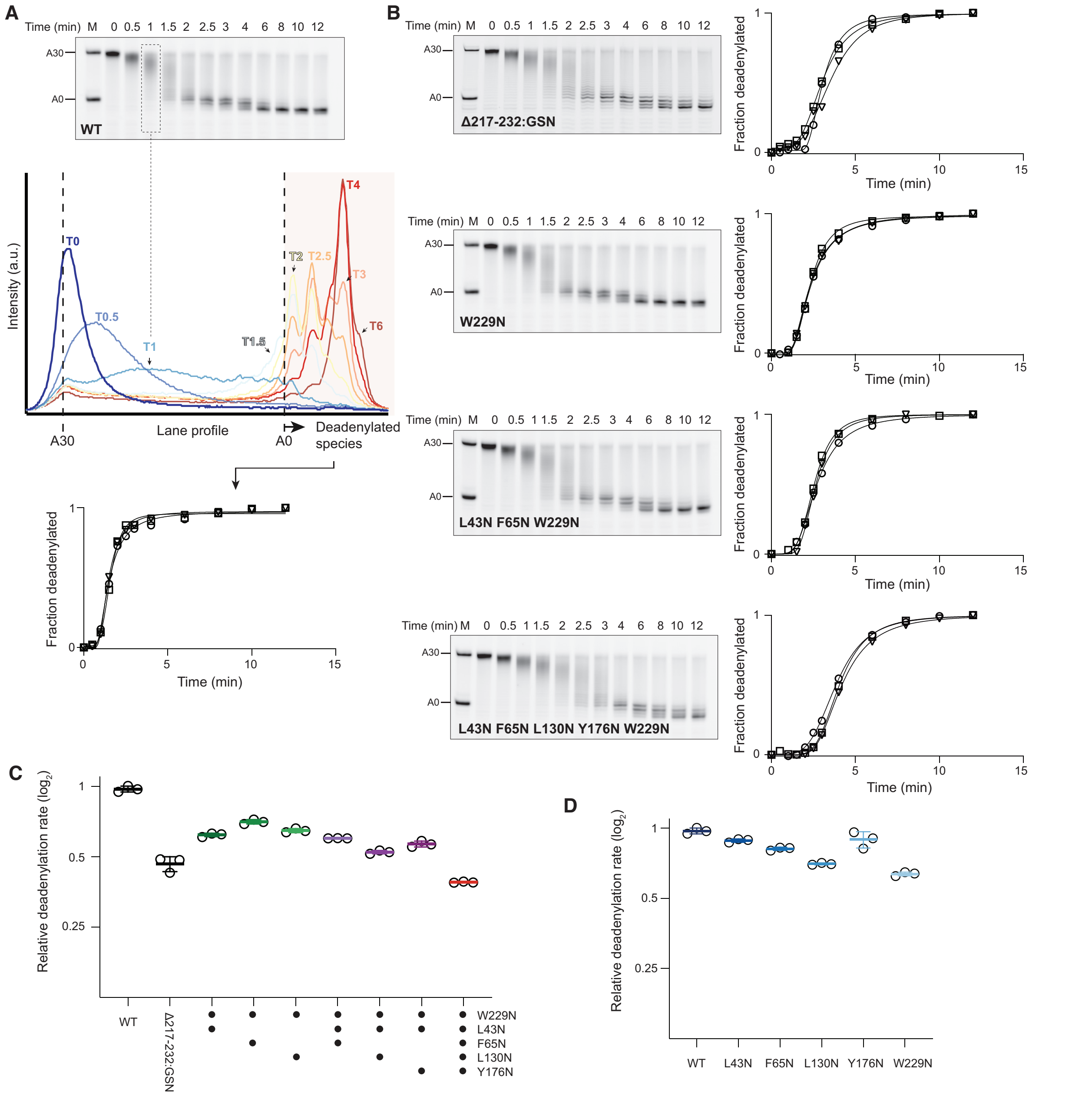


**Extended Data Fig. 5: Quantitation of deadenylation assays performed in the presence of Puf3 mutants.** (**A**) Representative deadenylation assay and densitometry analysis. Wild-type Puf3 (WT) was incubated to form a 1:1 complex with a 5' fluorescently labelled substrate RNA, containing a Pumilio response element (PRE) (UGUAAAUA) in a 20-nt region upstream of a 30-nt poly(A) tail. Puf3-bound RNAs were then incubated with recombinant Ccr4-Not complex and fractions were taken at indicated time points. Products were resolved on denaturing PAGE before imaging (top). Intensity profiles across each lane were integrated (middle) and the fraction of deadenylated RNA was quantified for each time point (bottom). Data were then fitted using an asymmetric logistic function to calculate the poly(A) tail half-life, enabling quantitation of relative deadenylation rates (see panel C). Assays were performed in triplicate with quantification shown for all replicates. M, molecular weight marker. (**B**) Representative assay gels and quantitation for mutant Puf3 proteins, as described in panel A. (**C,D**) Relative activities of Puf3 (C) combined point mutants and (D) single point mutants in stimulation of Ccr4-Not-mediated deadenylation. Assays and quantification were performed according to panel A and poly(A) tail half-lives are plotted relative to WT Puf3. Line, calculated mean from three independent replicates (open circles). Error bars represent the standard deviation.

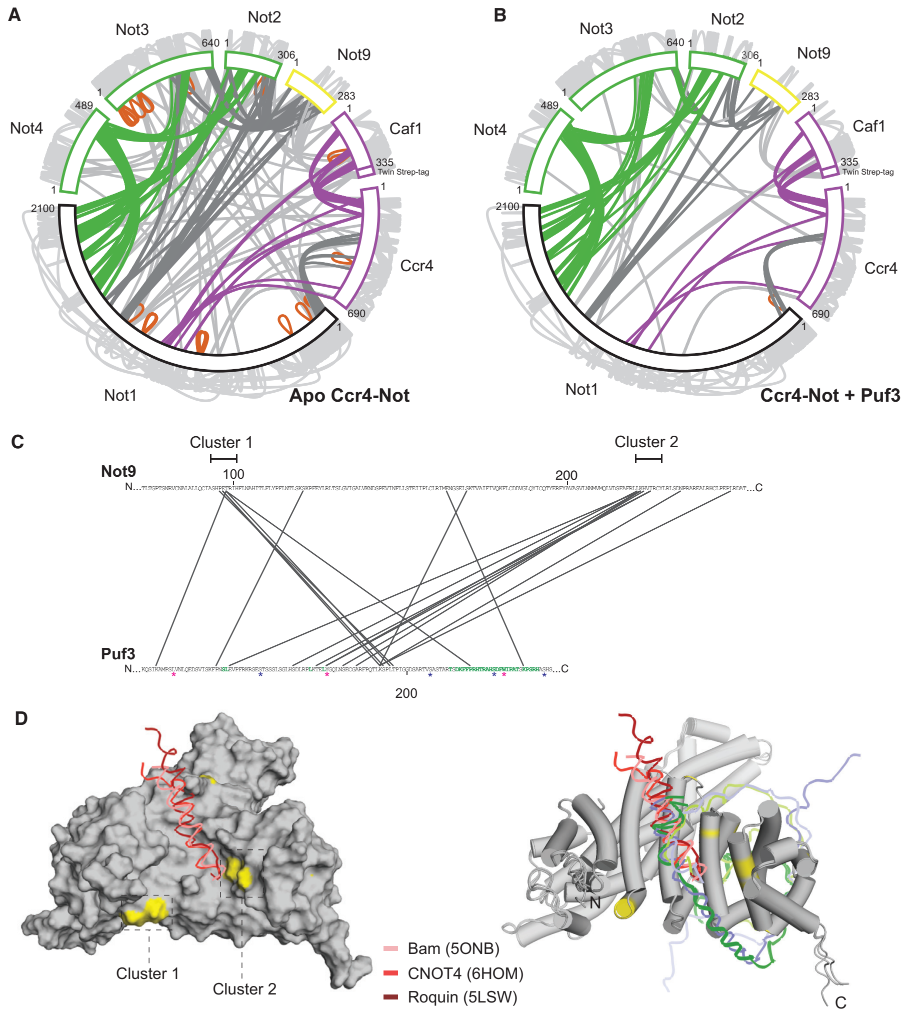


**Extended Data Fig. 6: Crosslinking mass spectrometry analysis of core Ccr4-Not subunits.** (**A-B**) Circular representation of crosslinking mass spectrometry analysis (CLMS) of *S. pombe* Ccr4-Not, alone (A) and in the presence of 1.5-fold molar excess of Puf3 (B). Recombinant full-length complexes were subjected to UV-activated Sulfo-SDA CLMS. Proteins and crosslinks are colored according to the subunit architecture of Ccr4-Not. Intra-protein crosslinks are displayed around the outside in grey. Self-links are shown in orange. To allow comparison, Puf3 crosslinks have been omitted in panel B. These data are consistent with no large scale conformation changes in Ccr4-Not on binding Puf3. (Also see Fig. 3A). (**C**) Detailed view of crosslinks between Not9 and Puf3 mapped onto segments of the protein sequences showing two distinct clusters on the Not9 subunit. Puf3 residues exchange-broadened in the presence of Ccr4-Not are colored green. Asterisks mark phosphorylated residues (blue) and residues mutated in this study (magenta). (**D**) Surface (left) and cartoon (right) representations of AlphaFold2 prediction of *S. pombe* Not9 and Not1. Peptides from Bam, CNOT4 and Roquin co-crystal structures with *H. sapiens* CNOT9 are overlaid. Puf3 CLMS clusters 1 and 2 around the peptide binding groove are highlighted in yellow. Two AlphaFold2 predictions of Puf3 binding to Not9 are also shown in the cartoon diagram (right) in purple and green.

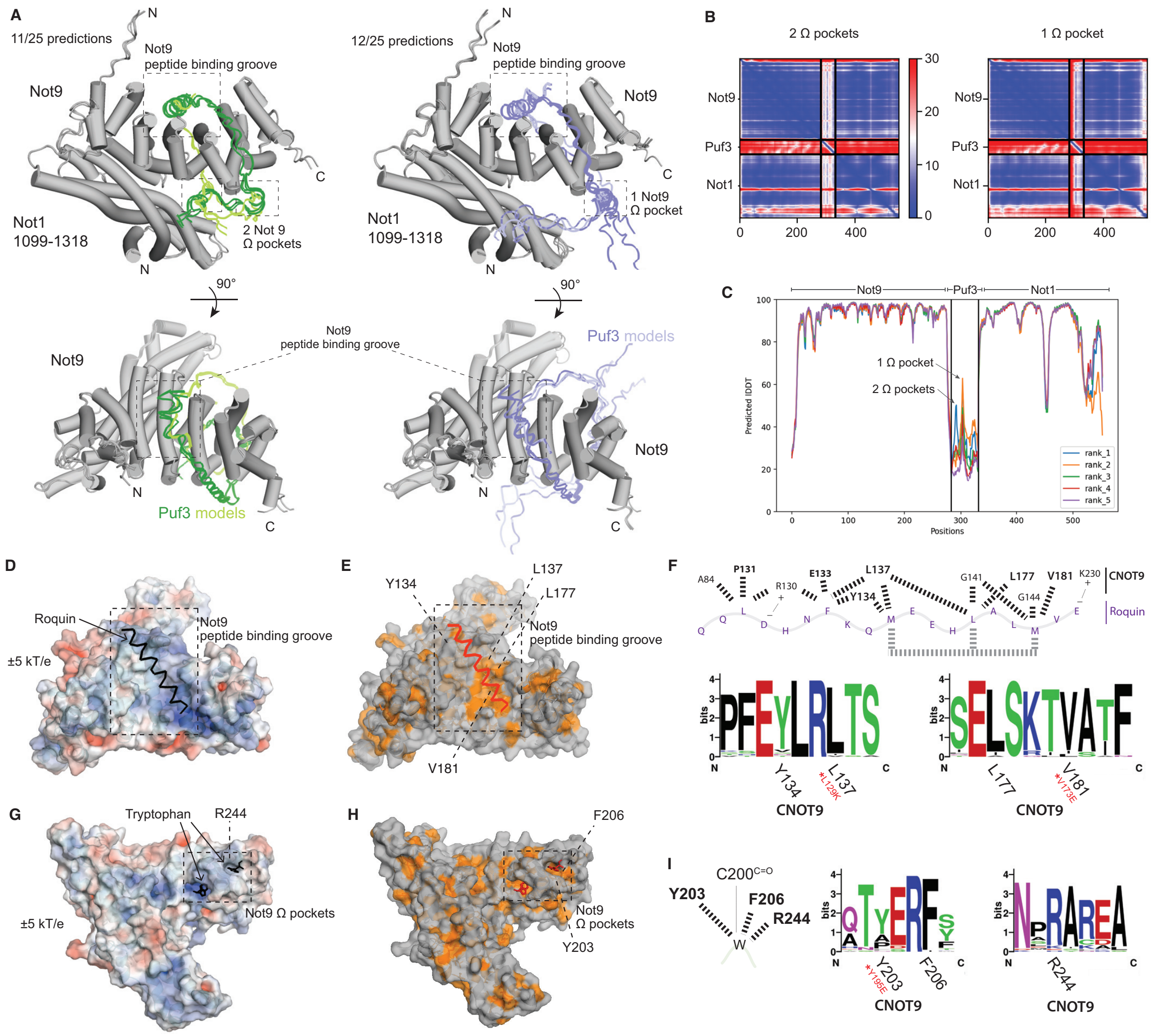


**Extended Data Fig. 7: Prediction and analysis of Puf3-Not9 interactions.** (**A**) Alphafold2 predicted structures for the complex formed between *S. pombe* Not9, Not1 residues 1099-1318 and Puf3 residues 209-265. 25 models were predicted from 5 runs, each with a single random seed and 5 models per run. Models were aligned to the structured domains of Not9 and Not1 and categorized according to whether one (right, blue ribbon) or two (left, green ribbon) of the previously identified Trp-binding pockets are engaged. These pockets can be more broadly defined as aromatic (Ω) pockets since they are also predicted to bind residues other than Trp (e.g Phe). Predicted structures are displayed in two orientations, showing that 23/25 predictions engage both the peptide binding groove and Ω -pocket(s).
(**B**) Representative PAE plots for predicted structures with one and two Ω-pockets in Not9 occupied (as shown in panel A). (**C**) Representative predicted lDDT scores per residue for 5 of the structural predictions shown in panel A. The confidence in Puf3 (middle) modelling is low but there is a local higher confidence in the region that binds the Ω-pockets of Not9. (**D-F**) Co-crystal structure of human CNOT9 bound to a peptide of Roquin (PDB 5LSW) shown as a surface representation colored by electrostatic surface potential (D) or by exposed hydrophobic residues (E) and as an interaction map (F), focusing on the peptide binding groove. In panel F, evolutionary conservation of CNOT9 (residues in bold in interaction map on top) is shown below in sequence logos generated using WebLogo from consensus sequence alignments representing a broad range of eukaryotic clades. Residues mutated in fission yeast Not9 in this study are marked with a red asterisk. (**G-I**) Co-crystal structure of human CNOT9 bound to Trp (PDB 4CRV) shown as a surface representation colored by electrostatic surface potential (G) or by exposed hydrophobic residues (H) and as an interaction map (I), all focusing on the Ω-pockets. Evolutionary conservation of CNOT9 residues is shown as described in panel F.

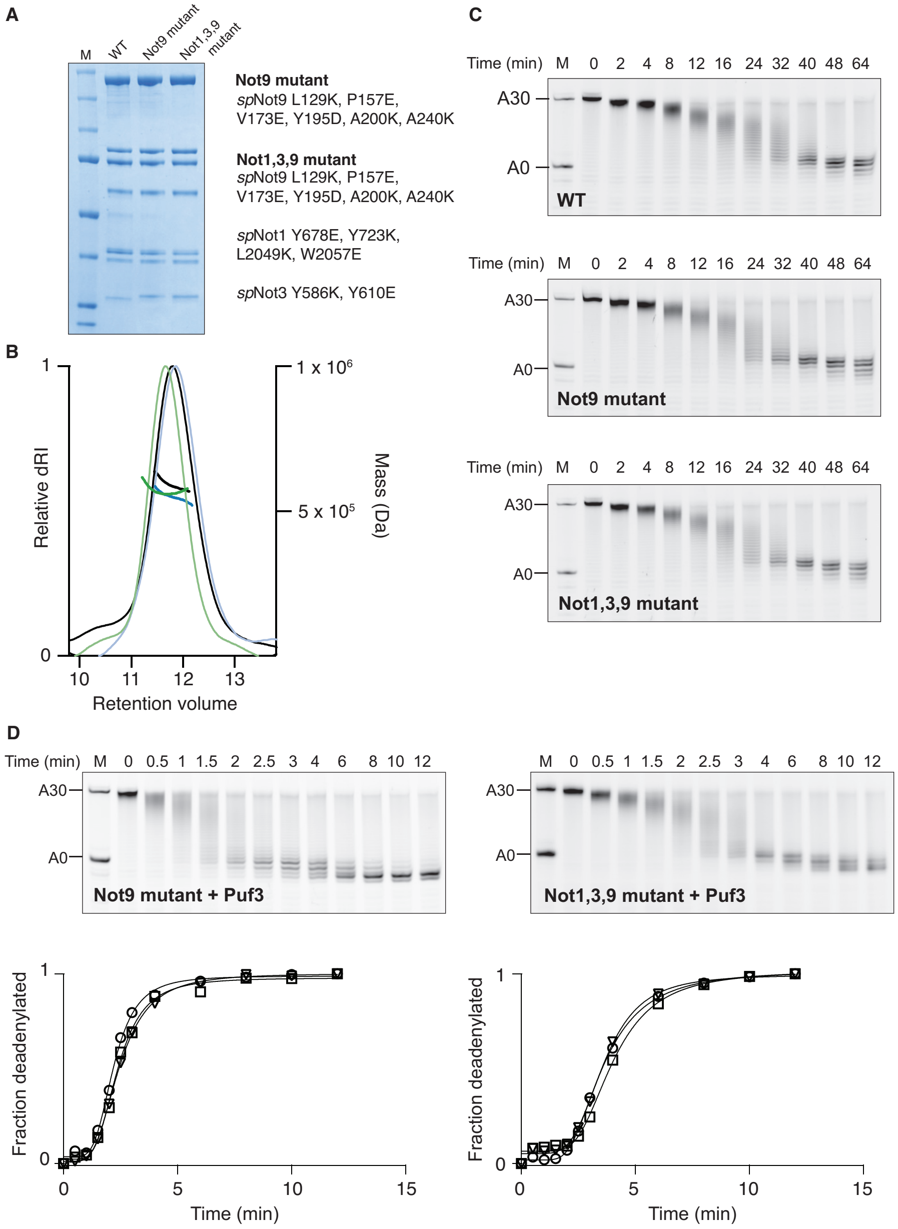


**Extended Data Fig. 8: Deadenylation activities of mutant Ccr4-Not complexes.** (**A**) Coomassie-stained SDS-PAGE analysis of purified recombinant Ccr4-Not complexes carrying indicated mutants in conserved SLiM binding sites, as detailed in Extended Data Fig. 7 and 9. M, molecular weight marker. (**B**) Size exclusion chromatography coupled to multiangle light scattering analysis of mutant complexes, showing that all complex variants are intact and monodisperse. 100 µl of each complex was injected on a Superose 6 increase 10/300gl column at 200 nM and analyzed using an inline coupled light scattering instrument. Peak traces (WT: Black, Not9 mutant: blue, Not1,3,9 mutant: green) show the normalized differential refractive index with the calculated mass plotted as lines (right axis). (**C**) Deadenylation assays of Ccr4-Not mutant complexes alone (without Puf3) on a substrate RNA containing a PRE and a 30-nt poly(A) tail, showing that there are no differences in intrinsic activity. (**D**) Representative deadenylation assay and corresponding densitometry analysis of indicated Ccr4-Not mutant complexes in the presence of WT Puf3 (related to Fig. 3B). Deadenylation assays and analyses were performed according to Extended Data Fig. 5A.

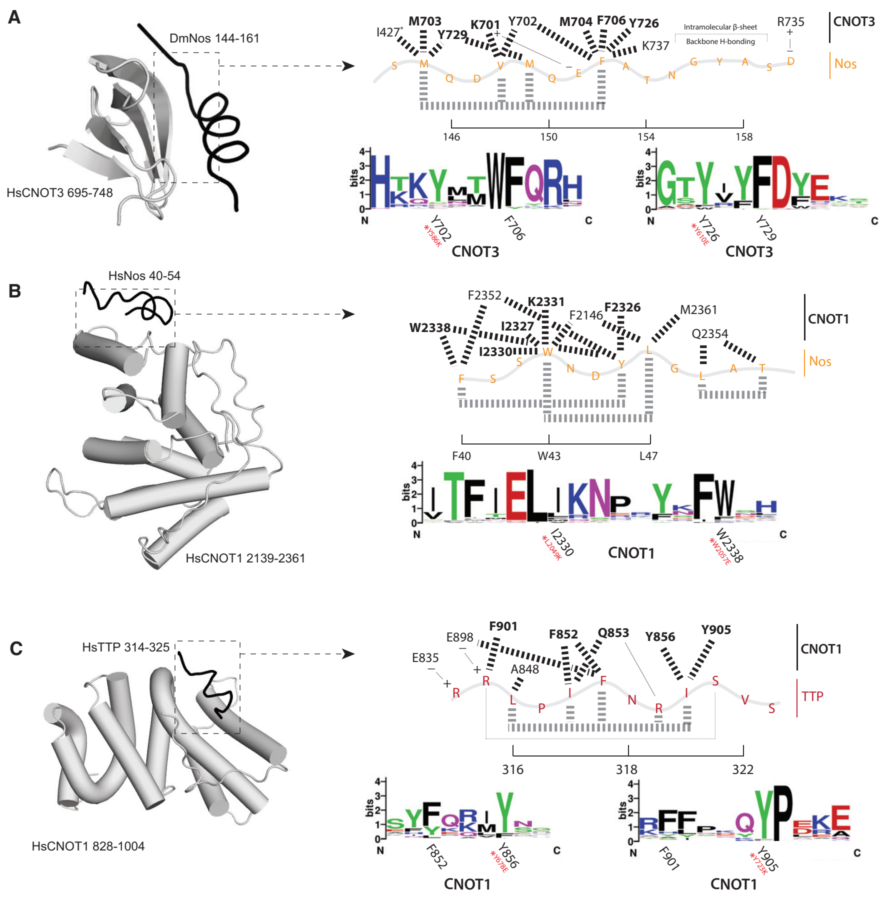


**Extended Data Fig. 9: Analysis of conserved CNOT1 and CNOT3 hydrophobic pockets.** (**A**) Analysis of a hydrophobic pocket in CNOT3. Left: CNOT3 centric view of the co-crystal structure between human (Hs) CNOT1, HsCNOT3 and *Drosophila* Nanos (DmNos) (PDB ID: 5FU7). Right: Diagrammatic representation of interactions between DmNos and HsCNOT3 and conservation analysis as in Extended Data Fig. 7F. Sites that were mutated in spNot3 are shown in red and marked with an asterisk. (**B**) Analysis of a hydrophobic pocket at the C-terminus of HsCNOT1 bound to HsNos (PDB ID: 4CQO), as in panel A. (**C**) Analysis of a hydrophobic pocket in the N-terminal HEAT repeat domain of HsCNOT1 bound to HsTTP (PDB ID: 4J8S), as in panel A.

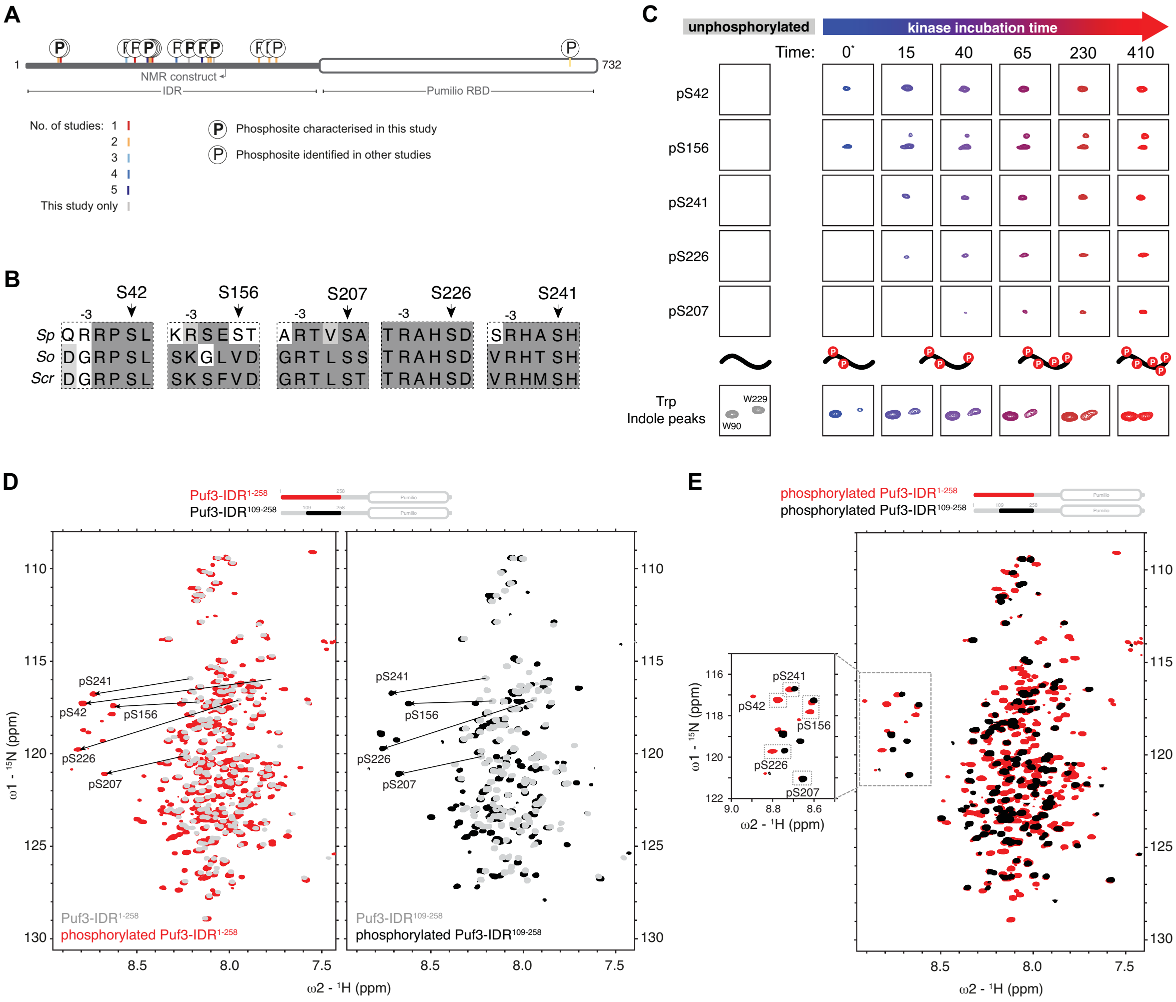


**Extended Data Fig. 10: Specific and ordered phosphorylation of the Puf3 IDR.** (**A**) Schematic of annotated Puf3 phosphorylation sites identified in this study and previous phosphoproteomics studies (as curated in PomBase). Color code shows the number of studies (including this one) identifying a particular site. Sites in bold are characterized using NMR spectroscopy in this study. (**B**) Alignments of *Schizo-saccharomycetes* Puf3 sequences around the identified phosphorylation sites. *Sp, S. pombe; So, S. octosporus; Scr, S. cryophilus*. Arg is often found in the -3 position. (**C**) Selected regions of 2D $^{1}H$ $^{15}N$ HSQC spectra of phosphorylation time courses. Addition of kinases leads to appearance of peaks in a distinct temporal pattern. The kinase was added at 0 minutes (asterisk), but the spectrum shows phosphorylation at S42 and S156 due to very rapid activity under these conditions and the time taken to load the sample and acquire the data. Trp indole peaks at the same time points are shown below reproduced from Fig. 4D. (**D**) Overlays of 2D $^{1}H$ $^{15}N$ HSQC spectra of non-phosphorylated and phosphorylated Puf3-IDR$^{1-258}$ and Puf3-IDR$^{109-258}$. (**E**) Overlay of 2D $^{1}H$ $^{15}N$ HSQC spectra of phosphorylated Puf3-IDR$^{1-258}$ and Puf3-IDR$^{109-258}$ samples used for assignment. Inset shows assigned peaks of phospho-serine residues.

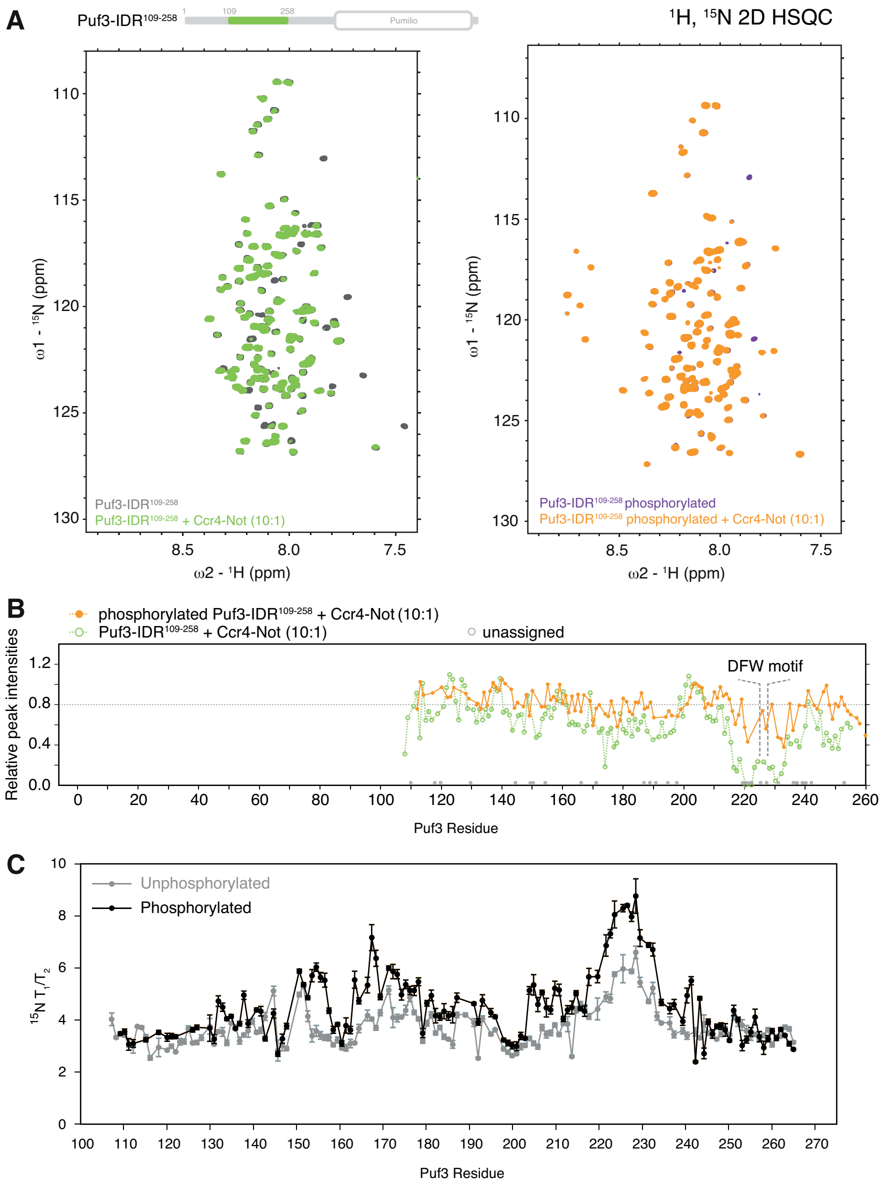


**Extended Data Fig. 11: Binding and dynamics of phosphorylated Puf3.** (**A**) $^{1}H,^{15}N$ 2D-HSQC of 75 μM Puf3-IDR$^{109\text{-}258}$. Spectra are shown for unphosphorylated (left) and phosphorylated (right) proteins with and without 7.5 μM unlabeled Ccr4-Not. (**B**) Relative peak intensities of non-phosphorylated (green open circles) and phosphorylated (yellow) Puf3-IDR$^{109\text{-}258}$ in the presence of Ccr4-Not from $^{1}H,^{15}N$ 2D-HSQC plotted from spectra in panel A. (**C**) $T_1$ and $T_2$ $^{15}N$-backbone dynamics of unphosphorylated and phosphorylated Puf3-IDR$^{109\text{-}258}$ used to generate difference plot in Fig 4E.

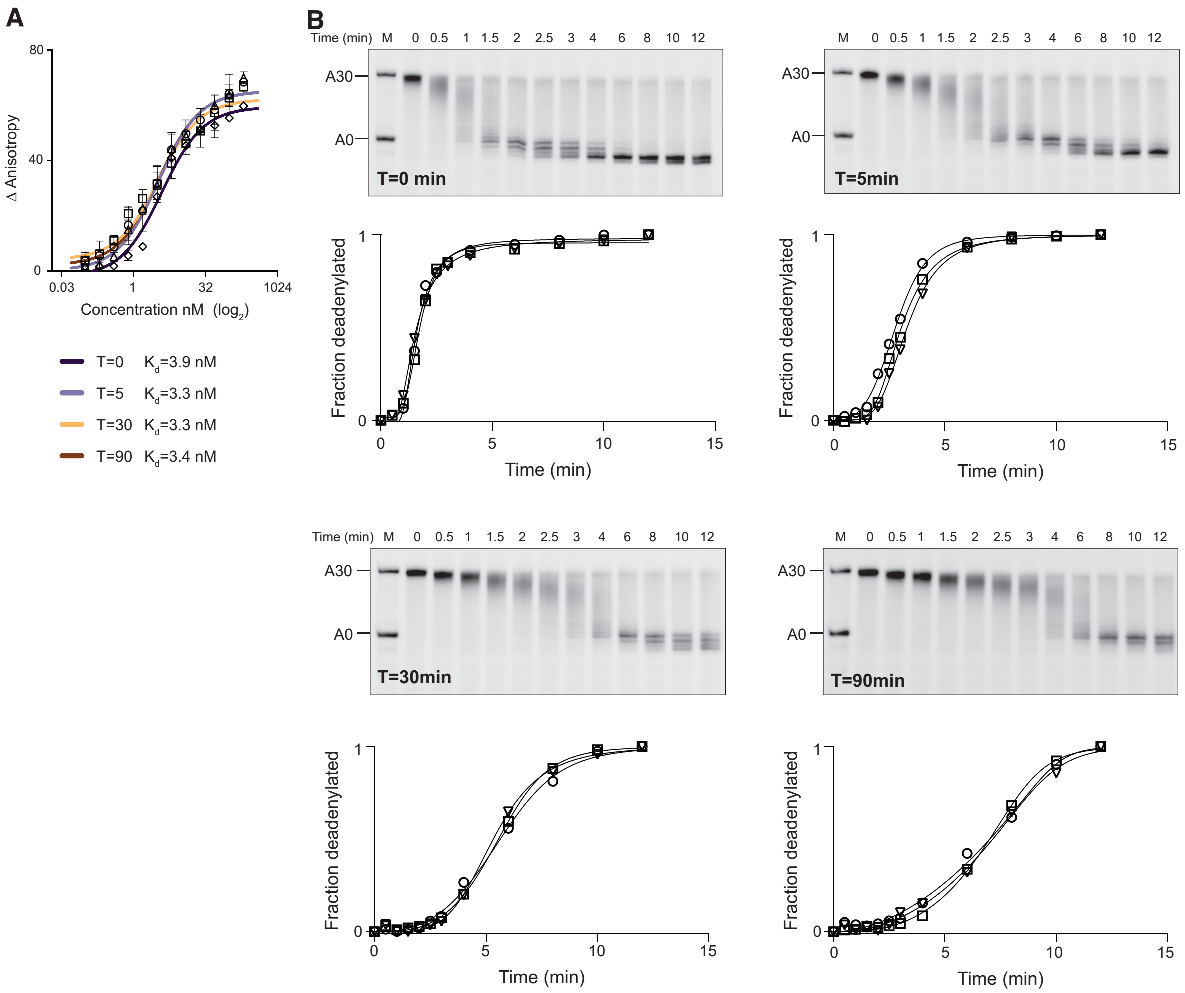


**Extended Data Fig. 12: RNA binding and deadenylation activity assays of phosphorylated Puf3.** (**A**) Fluorescence anisotropy of 5' FAM labelled PRE-containing RNA, titrated against increasing concentrations of MBP-Puf3 phosphorylated for the indicated time. Data were fitted with a quadratic binding equation to calculate the dissociation constants shown below. Data points are the mean of at least 3 technical replicates with error bars showing standard deviation. (**B**) Representative deadenylation assays and densitometry analysis of data plotted in Fig. 4F. The ability of full-length MBP-tagged Puf3 phosphorylated for the indicated times to stimulate the deadenylation activity of Ccr4-Not was analyzed according to Extended Data Fig. 5A. Three replicate assays are shown in each of the plots.

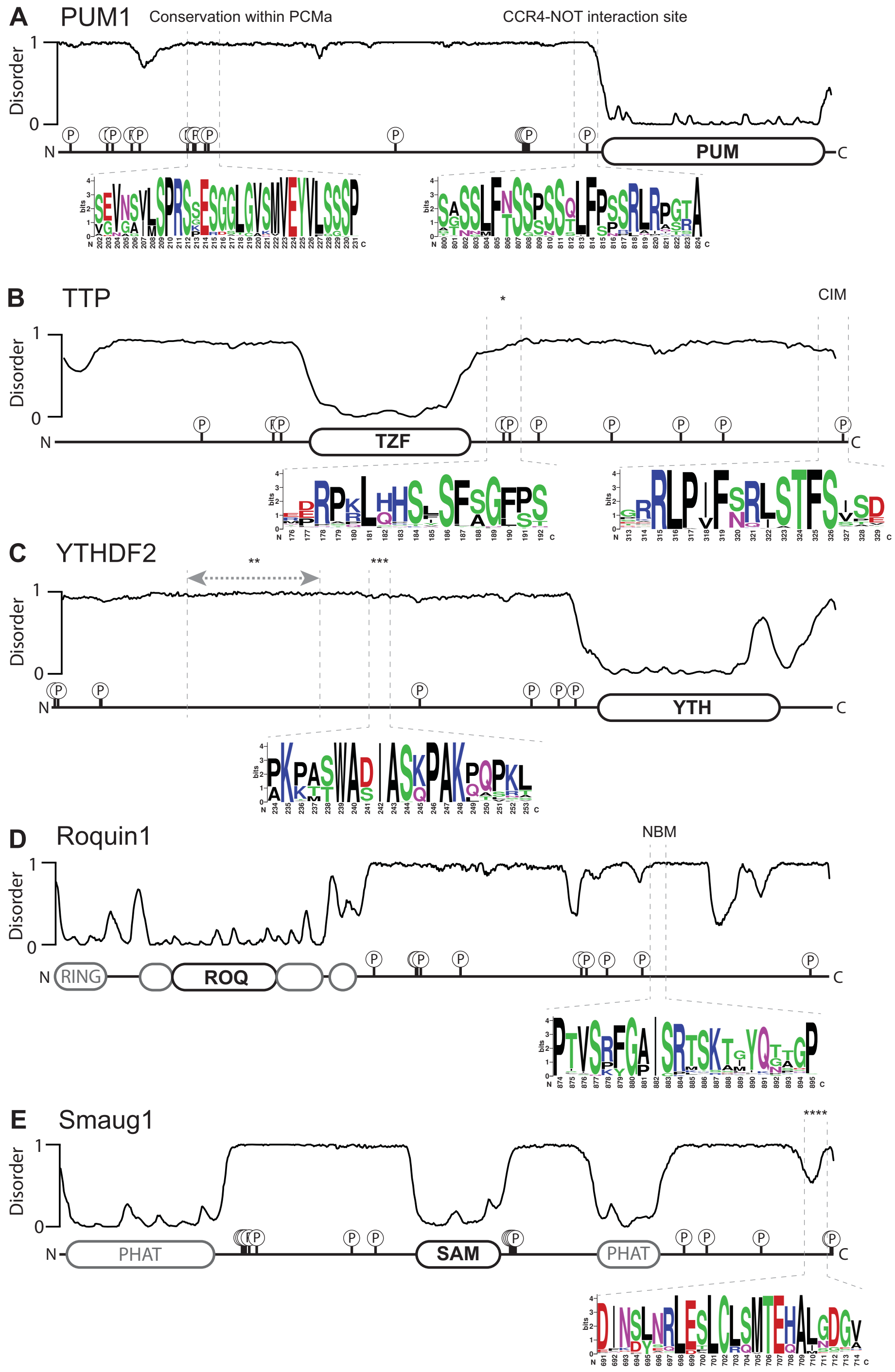


**Extended Data Fig. 13: Schematics of human CCR4-NOT interacting RNA-binding proteins.** Disorder prediction and domain diagrams of known *H. sapiens* RNA-binding proteins that recruit the CCR4-NOT complex. Structured domains are depicted as rounded rectangles, with known RNA-binding domains in bold, with IDRs as lines. IDR boundaries are defined based on metapredict[54]. Phosphorylation sites are annotated using data curated in PhosphoSitePlus [55]. Sequence logos show patches of conservation within IDRs which could be important for protein interactions and/or post-translational modifications, often containing conserved serines, aromatic and hydrophobic residues. Sequence logos were generated using Clustal omega alignments of 500-1000 homologous sequences from Chordates, Molluscs and Arthropods. (**A**) Pumilio homolog 1 (PUM1) contains a ~1000 amino acid IDR N-terminal to the Pumilio RNA-binding domain. Several regions within the IDR have been implicated in interaction with CCR4-NOT. Two conserved regions of interest have been identified in previous studies (sequence logos). The autonomously repressive Pumilio conserved motif a (PCMa) region contains several conserved phosphosites [17]. A separate region proximal to the RBD has been shown to interact with several subunits of the CCR4-NOT complex [16]. (**B**) Tristetraprolin (TTP) contains both N- and C-terminal IDRs relative to the RBD. Both have been shown to interact with CCR4-NOT. The major CCR4-NOT-interacting motif (CIM) lies close to the C-terminus of the protein [12]. Other regions within the C-terminal IDR contain both conserved aromatic and serine residues (which can be phosphorylated), indicated by an asterisk (*). (**C**) YTH domain-containing family protein 2 (YTHDF2) can directly bind the CCR4-NOT complex. A region of the N-terminal IDR spanning residues 100-200 (denoted by **) is sufficient for the interaction with the C-terminus of the NOT1 scaffold [56]. This region lies outside of the most conserved region in the IDR that contains a W-containing SLiM (shown in inset ***). This could be important for interaction with CCR4-NOT but equally could be involved in binding distinct protein partners or subcellular localization, e.g. via membrane binding. (**D**) Roquin contains multiple CCR4-NOT interacting motifs within its C-terminal IDR. Roquin has been shown to interact with both the NOT-module and CNOT9. Sequence logo shows conservation within the NOT-binding motif (NBM). (**E**) Smaug contains multiple IDRs with putative phosphoregulatory sites. Drosophila Smaug binds CCR4-NOT through an unknown interaction with the NOT module [57]. A C-terminal SLiM (****) is conserved and a possible candidate. This SLiM is within 10 amino acids of several C-terminal phosphosites (outside of sequence logo shown).

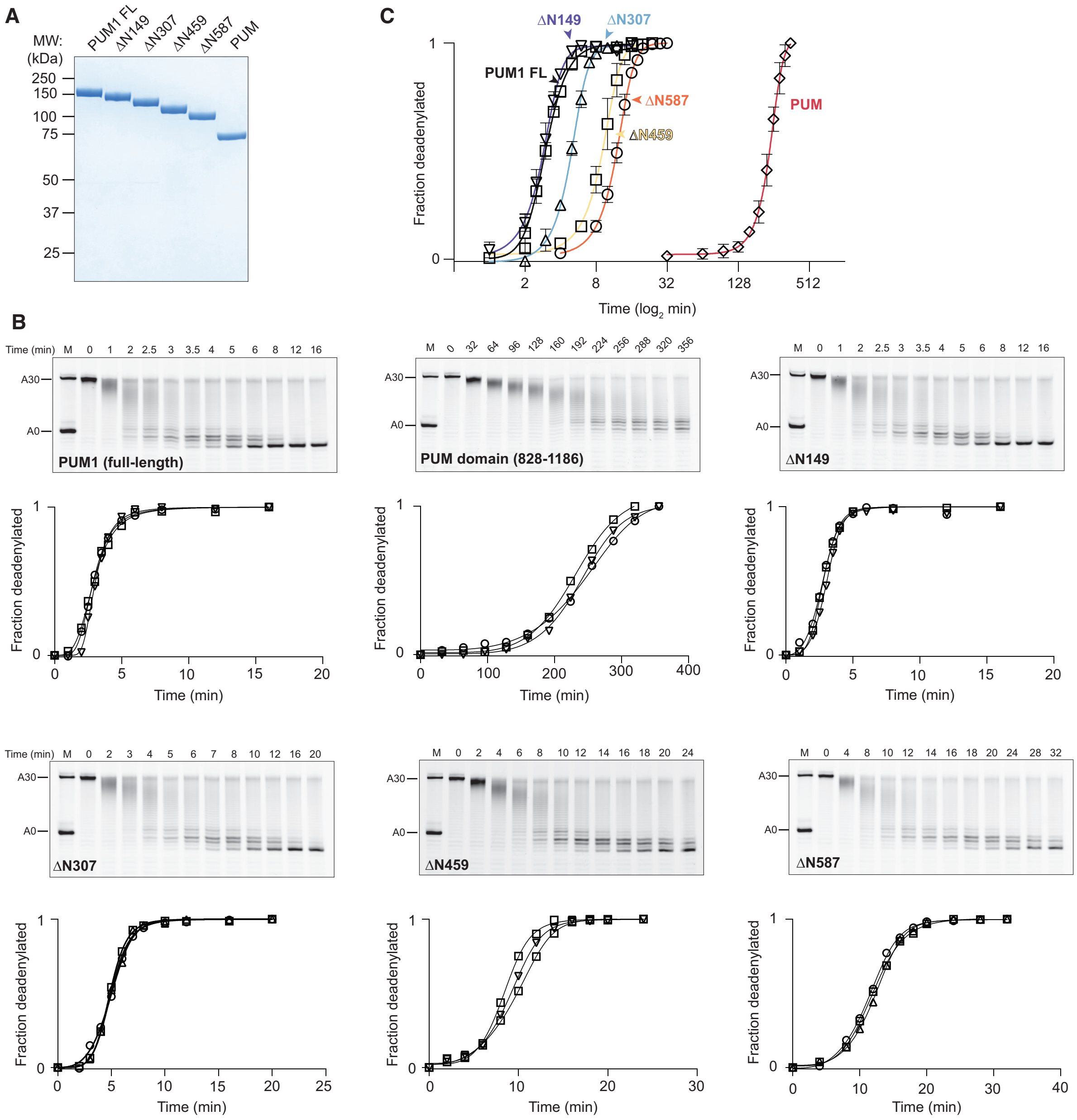

**Extended Data Fig. 14: Reconstitution of human Pumilio (PUM1) mediated targeted deadenylation.** (**A**) Purified PUM1 truncations. Truncations were designed based on previous studies defining conserved regions of the IDR. PUM, Pumilio domain alone. (**B**) *In vitro* deadenylation assays probing the ability of recombinant MBP-tagged HsPUM1 protein constructs to accelerate the exonucleolytic shortening of poly(A) tails from a synthetic RNA substrate by full-length recombinant human CCR4-NOT complex. MBP-tagged PUM1 constructs were preincubated to form a 1:1 complex with a 5' 6-FAM labelled substrate RNA containing a Pumilio response element (PRE) (UGUAAAUA) in a 20-nt upstream region and a 30-nt poly(A) tail. Reactions were started by the addition of 50 nM CCR4-NOT complex, samples withdrawn at indicated time points, and reaction species separated on denaturing PAGE gels before imaging. One representative gel is shown for each construct and three replicates are shown in each plot. (**C**) Quantification of deadenylation assays (from panel B). Open shapes for each construct show average of 3 replicate assays. Error bars show standard deviation. The Δ587 construct containing ~250 predicted unstructured residues proximal to the Pumilio RNA-binding domain retains the ability to stimulate removal of the poly(A) tail by ~10-fold.

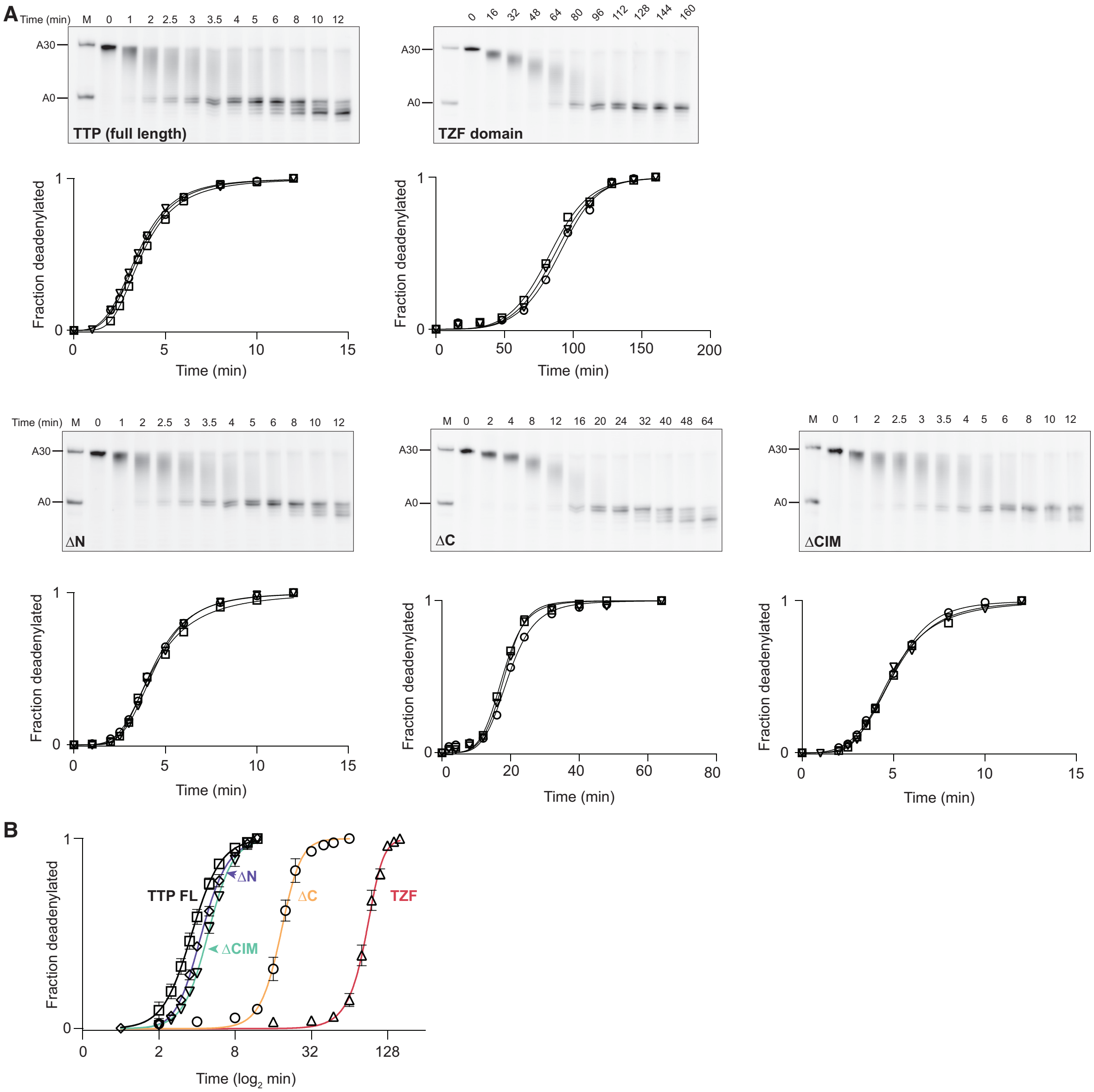


**Extended Data Fig. 15: Reconstitution of targeted deadenylation with human Tristetraprolin (TTP).** (**A**) *In vitro* deadenylation assays testing the ability of recombinant human MBP-tagged TTP protein IDR deletion constructs to accelerate the deadenylation activity of full-length recombinant *H. sapiens* CCR4-NOT complex. Indicated MBP-tagged TTP constructs were preincubated to form a 1:1 complex with a 5' 6-FAM labelled substrate RNA containing an AU-rich element (ARE) (UUAUUUAUU) in a 20-nt upstream region and a 30-nt poly(A) tail. Reactions were started by the addition of 50 nM CCR4-NOT complex and samples withdrawn at indicated time points before reaction species were separated on denaturing PAGE gels and imaged. A representative gel is shown for each construct and all three replicates are plotted. (**B**) Quantification of deadenylation assays for constructs shown in panel A. Open shapes for each construct show the average of 3 replicate assays with error bars showing standard deviation. The structurally characterized interaction between the CIM motif and NOT1 subunit is not sufficient to fully account for the stimulatory activity of TTP, consistent with previous data [13].

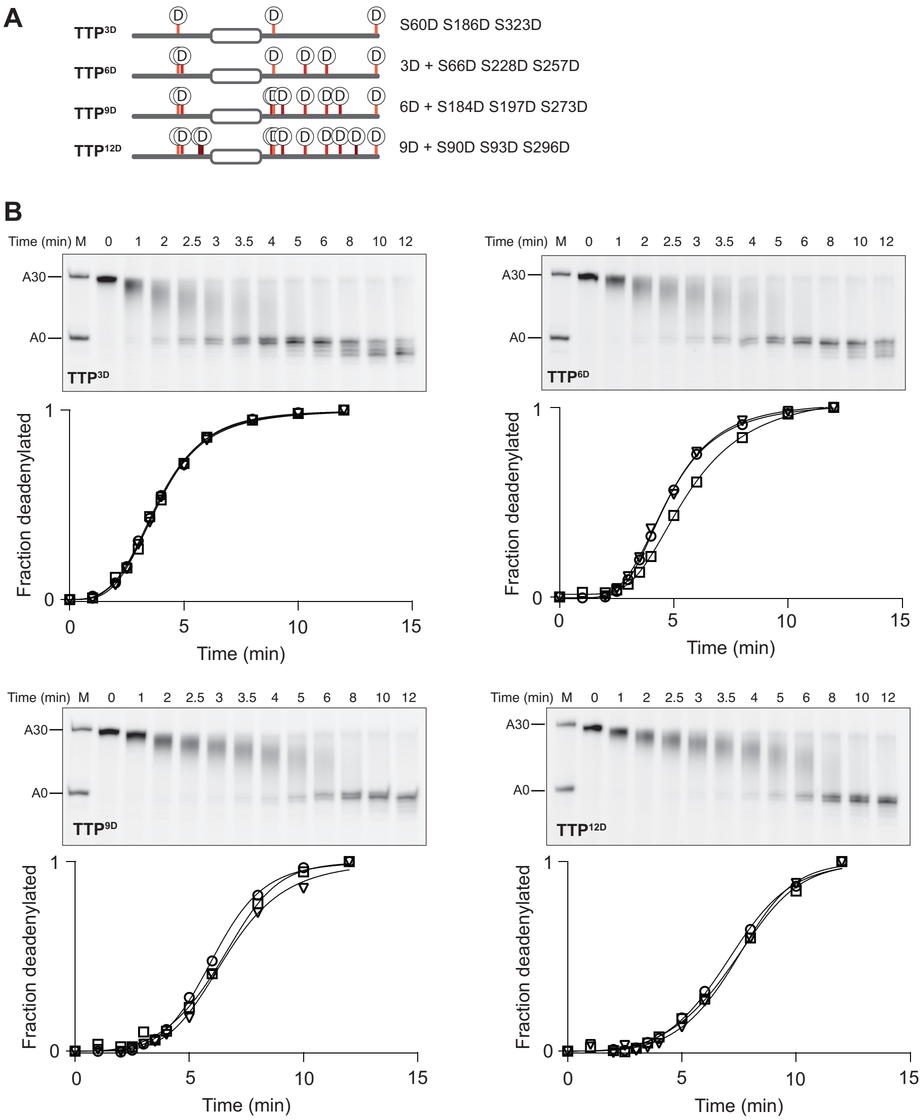


**Extended Data Fig. 16: Deadenylation in the presence of phosphomimetic TTP proteins.** (**A**) Phosphomimetic TTP constructs used in this study. All phosphorylation sites have previously been characterized using either phosphoproteomics MS or phosphopeptide mapping. Point mutant proteins at indicated positions (mutated to Asp) were cloned, overexpressed and purified to homogeneity for assays performed in panel B. (**B**) *In vitro* deadenylation assays performed as in Extended Data Fig. 15 but with indicated TTP phosphomimetic constructs.
